# TRiC folds the giant ciliary protein IFT172 via a non-canonical open-state mechanism

**DOI:** 10.64898/2026.03.28.714460

**Authors:** Qiaoyu Zhao, Jiayi Li, Yujie Tong, Yurui Li, Wenyu Han, Zuyang Li, Yanxing Wang, Yue Yin, Jun Fang, Wanying Jiang, Qianqian Song, Shengyou Huang, Yidong Shen, Yao Cong

## Abstract

The eukaryotic chaperonin TRiC/CCT is essential for folding complex proteins, yet how it folds substrates that exceed its closed chamber capacity remains a longstanding paradox. Here, we define the folding pathway of IFT172, the largest subunit (∼200 kDa) of the intraflagellar transport (IFT) machinery, and uncover a “divide-and-conquer” mechanism. TRiC and HSP70 engage IFT172 concurrently but on distinct domains: TRiC captures the N-terminal WD40 β-propellers within its chamber, whereas HSP70 independently stabilizes the C-terminal TPR domain in the cytosol. To accommodate this oversized client, specific TRiC subunits (CCT4, CCT2, and CCT7) undergo pronounced Z-shaped outward bending, thereby expanding the chamber. Unexpectedly, the first WD40 domain reaches a near-folded state within the open, ATP-bound chamber, and subsequent TRiC ring closure triggers substrate ejection rather than encapsulation. This non-canonical “fold-and-eject” mechanism challenges the classical view that the closed chamber is an obligate folding cage. We further demonstrate that this pathway is essential for ciliary functions *in vivo,* and reveal a conserved mode of chaperonin recognition among IFT components bearing tandem WD40-TPR architectures. Together, our findings establish a new paradigm for the folding of oversized, multi-domain proteins and identify TRiC as a central proteostasis hub in ciliary biogenesis, with direct implications for ciliopathy pathogenesis.

## Introduction

Eukaryotic proteomes are enriched in large, multi-domain proteins whose folding demands exceed intrinsic capacity and therefore require assistance from molecular chaperones^1,2^. The group II chaperonin TRiC/CCT occupies a central position in this network, mediating the folding of ∼10% of the cytosolic proteome, including cytoskeletal proteins, cell-cycle regulators^3,4,5,6^, and an expanding class of WD40 β-propeller proteins such as CDC20, Gβ, CSA, TAF and mLST8^7–11^. The canonical mechanism of TRiC-assisted folding, established through decades of studies on relatively small substrates (<55 kDa), invokes an ATP-driven conformational cycle in which ring closure encapsulates the substrate within a shielded cage, where steric confinement and a favorable chemical environment promote productive folding^12–17^. This encapsulation model has been reinforced by recent cryo-EM structures of TRiC bound to actin, tubulin, and Gβ, all of which fold within the closed chamber^13–17^.

Yet, TRiC is also required for the biogenesis of proteins that far exceed the ∼70 kDa volumetric capacity of its closed cavity^18^. How the chaperonin accommodates and productively folds such oversized, multi-domain clients, which cannot be fully encapsulated, remains a fundamental unresolved question. Although partial encapsulation has been proposed as a possible solution^18^, direct structural visualization of how TRiC accommodates oversized clients, whether it coordinates with other chaperone systems during this process, and at which stage of the ATPase cycle folding actually occurs has been lacking.

This knowledge gap is particularly consequential for the biogenesis of intraflagellar transport (IFT) complexes, the molecular machines that build and maintain cilia^19–21^. IFT complexes, consisting of IFT-A (6 subunits) and IFT-B (16 subunits) subcomplex, comprise numerous large, multi-domain subunits whose precise folding is essential for ciliary function. Accordingly, mutations that compromise the folding or stability of IFT components are a primary cause of ciliopathies, a class of genetic disorders affecting nearly every organ system^22,23^. IFT172 exemplifies this challenge: as the largest subunit (∼200 kDa) of IFT-B, it contains two tandem N-terminal WD40 β-propeller domains followed by an extended C-terminal TPR solenoid, an architecture that far exceeds the capacity of the TRiC folding chamber. How cells ensure the efficient folding of IFT172 and its assembly into functional IFT machinery, and whether defects in this process contribute to ciliopathy pathogenesis, remain critical unanswered questions.

Here, we integrate cryo-EM, cross-linking mass spectrometry (XL-MS), biochemical analysis, and *in vivo* functional assays to define the TRiC-assisted folding pathway of IFT172. We uncover a “divide-and-conquer” strategy in which TRiC and HSP70 simultaneously engage distinct structural domains of IFT172, coupled with allosteric remodeling of the TRiC chamber to accommodate this oversized substrate. Unexpectedly, our cryo-EM structures reveal that the WD40-1 domain reaches a near-folded conformation within the open, ATP-bound state of TRiC, before ring closure, challenging the longstanding view that encapsulation is obligatory for productive folding. We further show that this pathway is essential for ciliary function *in vivo*, and reveal a conserved mode of chaperonin recognition among IFT subunits sharing the tandem WD40-TPR topology. Together, these findings redefine the mechanistic logic of TRiC-assisted folding, establish TRiC as a central proteostasis hub for ciliary biogenesis, and illuminate the evolutionary sophistication by which the chaperonin system has adapted to the demands of the complex eukaryotic proteome.

## Results

### TRiC participates in the biogenesis of IFT172

To identify chaperones involved in IFT172 biogenesis, we overexpressed FLAG-tagged IFT172 in HEK293F cells, affinity-purified the protein, and characterized its interactome by mass spectrometry. In addition to 9 of other 15 IFT-B subunits, we detected strong enrichment of molecular chaperones, including HSP70, Hsp90, and all eight TRiC subunits (Fig. 1A). Co-immunoprecipitation and glycerol gradient centrifugation confirmed that IFT172 and TRiC form a stable complex (Fig. 1B, C, Fig. S1A-B). Native-gel migration and SDS-PAGE further showed co-migration of IFT172 with TRiC (Fig. 1D-E). Mass spectrometry of the TRiC band excised from the native gel also identified IFT172 together with HSP70, as well as low levels of other TRiC clients, including tubulin and actin (Fig. S1C, Table. S2). Notably, the TRiC-IFT172 complex retained ATPase activity comparable to that of TRiC alone (Fig. S1D), indicating that client engagement does not allosterically inhibit nucleotide hydrolysis ability of the chaperonin. All together, these results establish that IFT172 forms a stable complex with TRiC and HSP70, implicating a cooperative chaperone network in its biogenesis.

**Fig. 1.**
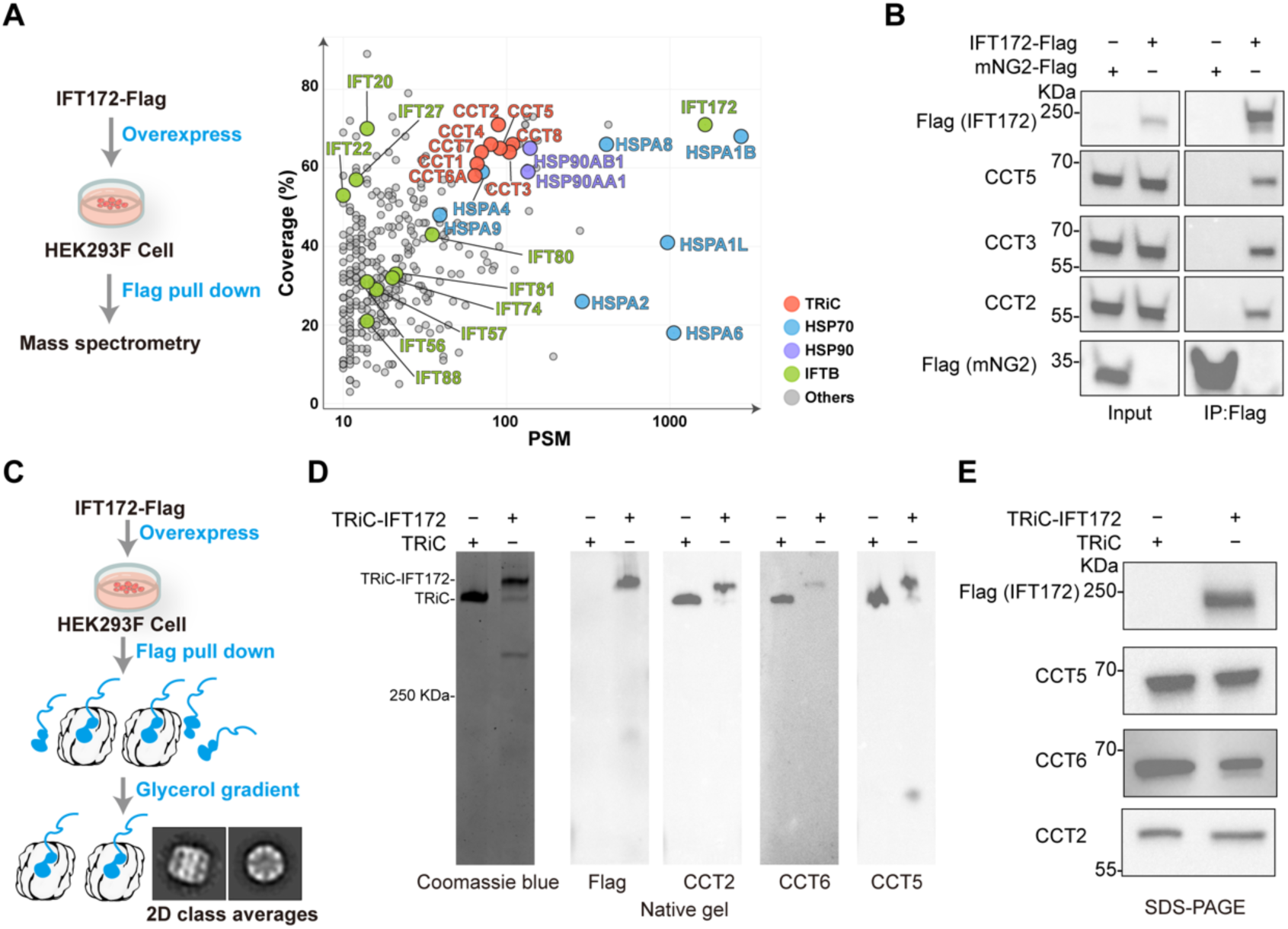
IFT172 forms a complex with TRiC. (A) FLAG-IFT172 affinity purification followed by mass spectrometry identifies TRiC and a network of chaperones as enriched interactors. (B) Co-immunoprecipitation from HEK293T cells expressing FLAG-IFT172 confirms association with endogenous TRiC subunits; FLAG-mNG2 was used as a negative control. (C) Schematic of the purification workflow used to isolate the TRiC-IFT172 complex, with reference-free 2D class averages of NS-EM data showing features of co-purified TRiC. (D) Native gel and immunoblotting show co-migration of IFT172 with TRiC, supporting formation of a stable complex. (E) Components analysis of the purified material by SDS–PAGE identifies IFT172, and TRiC subunits.

### Cryo-EM structure of the TRiC-IFT172 complex

To define how TRiC engages this oversized, multi-domain substrate, we determined the cryo-EM structure of the cross-linked TRiC-IFT172 complex at 3.55 Å resolution (Fig. 2A, Fig. S2A-D). The reconstruction revealed TRiC in an open, apo state with unoccupied nucleotide-binding pockets (Fig. 2A, Fig. S2E). Additional density within the TRiC chamber, distinct from the disordered N- and C-terminal tails of TRiC subunits, was attributable to substrate (Fig. 2A, Fig. S3A). Although the intrinsic flexibility of bound IFT172 limited local resolution, this density was clearly distinct in size and morphology from those of previously characterized TRiC-bound substrates (actin, tubulin and Gβ) and cofactors (PDCD5, prefoldin, or PHLP2)^14–17,24,25^. The substrate density was positioned near the CCT3/6/8/7 subunits and contacted the apical (A) and equatorial (E) domains of CCT3, CCT6, CCT8, CCT7, and CCT4, together with extensive contacts to the disordered N- and C-terminal tails of TRiC subunits, suggesting an important role for these flexible tail regions in IFT172 engagement (Fig. 2A, Fig. S3A). Notably, the density extended from the chamber interior through the apical opening to the exterior, consistent with a topology in which part of IFT172 protrudes beyond the TRiC cavity.

**Fig. 2.**
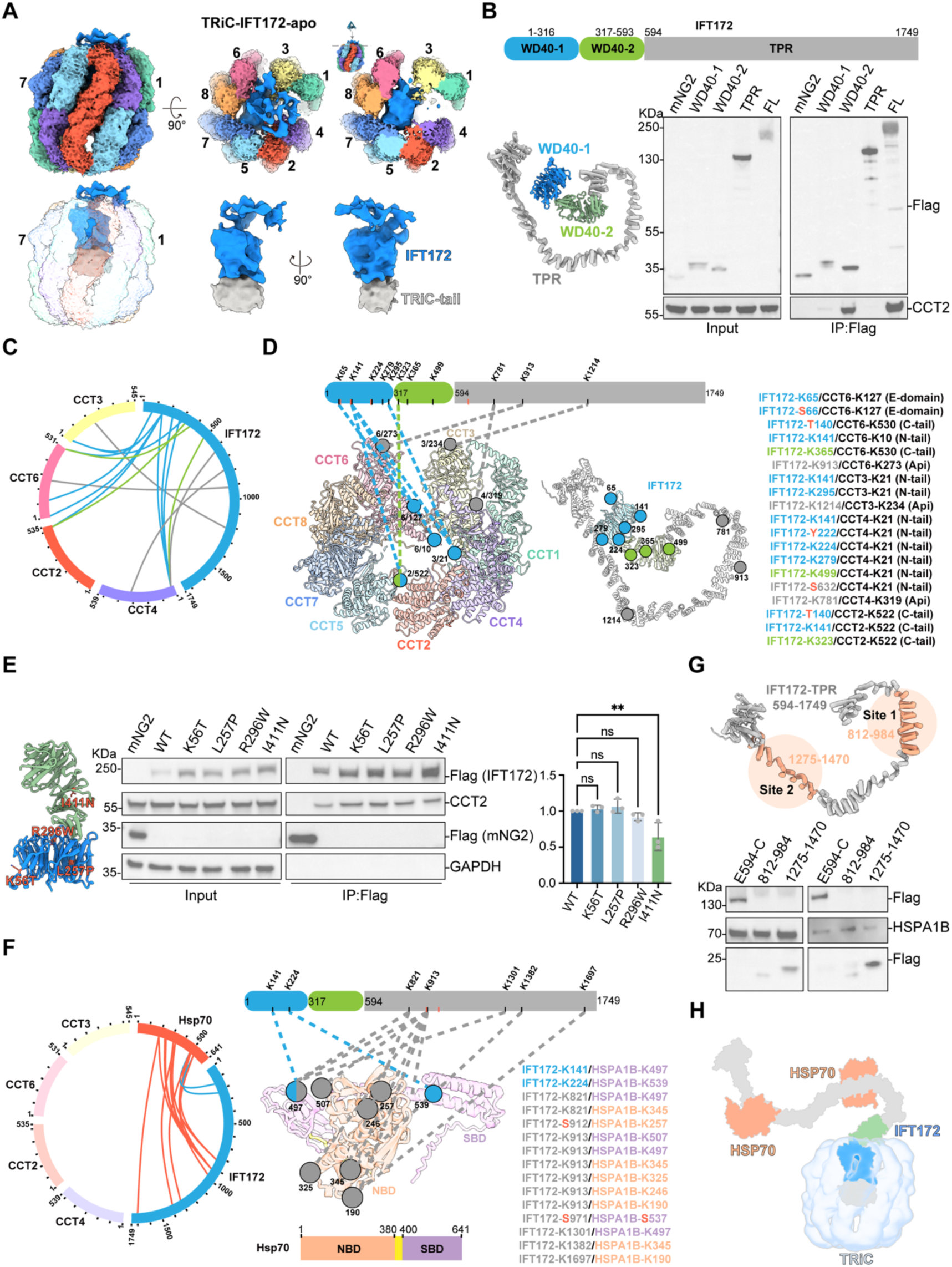
TRiC and HSP70 engage distinct domains of IFT172 to coordinate folding. (A) Cryo-EM reconstruction of the TRiC-IFT172 complex reveals additional density within the TRiC chamber. In the right panel of the first row, the fuzzy density atop TRiC has been clipped to visualize the substate interactions with the TRiC subunits’ apical domain. (B) Co-immunoprecipitation of transiently expressed IFT172 truncations (WD40-1, blue; WD40-2, green; and TPR domain, gray) from HEK293T cells shows that TRiC subunit CCT2 binds the WD40-1 and WD40-2 domains but not the isolated TPR domain (n = 3). This IFT172 color scheme is used throughout. (C) Crosslinking mass spectrometry (XL-MS) analysis of the TRiC-IFT172 complex. Circular plot summarizing inter-subunit crosslinks between IFT172 and TRiC subunits. Crosslinks are colored by IFT172 domain. (D) Mapping of crosslinked residues onto the AlphaFold3-predicted IFT172 model and the apo-state TRiC structure. Interactions involving unstructured TRiC N-/C-terminal tails are excluded. All 19 crosslinks are listed on the right. Crosslinks involving non-lysine residues (T/Y/S) that are supported by spectra are indicated in red. (E) Co-immunoprecipitation of wild-type and ciliopathy-associated IFT172 mutants transiently expressed in HEK293T cells, probed for TRiC binding. Right, quantification of CCT2 co-precipitation normalized to wild-type (mean ± SEM; n = 3). (F) XL-MS analysis reveals HSP70-IFT172 interactions. Left, circular plot summarizing crosslinks between HSP70 and IFT172; no crosslinks were detected between HSP70 and TRiC. Right, mapping of IFT172-HSP70 crosslinks onto the AlphaFold3-predicted model of HSP70 (NBD, salmon; SBD, light purple). (G) Co-immunoprecipitation of IFT172 TPR truncations identifies two HSP70-binding regions (Site 1, residues 812–984; Site 2, residues 1275–1470) and confirms binding to the full-length TPR domain (594–C terminus) (n = 3). (H) Proposed model for domain-partitioned chaperone action: TRiC engages the N-terminal WD40 domains of IFT172 within its chamber, whereas HSP70 binds two sites in the C-terminal TPR domain solenoid extending outside the chamber.

### TRiC recognizes and engages the WD40 domains of IFT172

To define the TRiC-binding region of IFT172, we expressed IFT172 truncations corresponding to WD40-1 (residue 1-316), WD40-2 (residue 317-593), and the TPR domain (residue 594-1749). Co-immunoprecipitation showed TRiC association with both WD40 domains, whereas the isolated TPR domain showed no detectable binding (Fig. 2B), identifying the N-terminal WD40 domains as the primary recognition elements for TRiC engagement.

We next performed XL-MS with BS^3^ as crosslinker to map the binding interface. Crosslinking stabilized the TRiC-IFT172 complex, as confirmed by SDS-PAGE and by the appearance of a faster-migrating species on native gel (Fig. S4A-B). As an internal quality control, we identified 54 inter-subunit crosslinks within TRiC, 92.6% of which satisfied the expected Cα-Cα distance threshold of 35 Å^26,27^ (Fig. S4C-E). Consistent with the truncation data (Fig. 2B), 15 of the 19 IFT172-TRiC crosslinks mapped to the WD40 domains (Fig. 2C), with extensive contacts involving the disordered terminal tails of CCT2, CCT3, CCT4, and CCT6 subunits. The remaining crosslinks in the TPR domain (K781, K913, and K1214) mapped to the A-domains of CCT3, CCT4, and CCT6 (Fig. 2D, Table. S3). Because the isolated TPR domain does not bind TRiC (Fig. 2B), these contacts likely reflect transient interactions made by the protruding TPR segment rather than primary recognition events.

To assess the folding status of the bound substrate, we mapped 13 intra-molecular crosslinks within the WD40 domains onto the folded AlphaFold-predicted model of IFT172. Only 53.8% satisfied the 35 Å permissible Cα-Cα distance restraint (Fig. S4F-H), indicating that the WD40 domains adopt a non-native or partially folded conformation upon initially bound to TRiC. Together, the cryo-EM and XL-MS data support a model in which TRiC captures the N-terminal WD40 domains of IFT172 within its chamber primarily through contacts with the disordered N- and C-terminal tails of the E-domains, while the C-terminal TPR solenoid extends through the apical opening of TRiC into the cytosol.

To link TRiC recognition to disease, we examined ciliopathy-associated IFT172 mutations that map to the WD40 domains, including K56T (cholestatic liver disease)^28^, L257P (Bardet-Biedl syndrome, BBS)^22^, and I411N/R296W (Jeune and Mainzer-Saldino Syndromes)^29^. The severe I411N mutation^29^, located in WD40-2, markedly reduced TRiC binding compared to wild-type (WT) IFT172, whereas K56T, L257P, and R296W retained near-normal association (Fig. 2E), implicating defective chaperonin recognition as a mechanism underlying impaired IFT172 biogenesis in ciliopathies.

### HSP70 chaperones the cytosolic exposed TPR domain of IFT172

The co-purification of HSPA1B (an HSP70 family chaperone, hereafter referred to as HSP70) with the TRiC-IFT172 complex suggested a collaborative role for this chaperone in IFT172 biogenesis (Fig. S1C). To define the structural basis of this cooperation, we mapped the HSP70-IFT172 interaction by XL-MS. The crosslinking profile revealed a striking domain specificity: only 2 crosslinks mapped to the N-terminal WD40 domains, whereas 13 mapped across the C-terminal TPR solenoid, indicating that HSP70 predominantly engages the TPR region (Fig. 2F, Table. S3). Notably, no direct crosslinks were detected between HSP70 and TRiC, indicating that the two chaperone systems engage IFT172 simultaneously yet independently on distinct domains, without forming a stable direct complex.

Mapping the crosslinked sites onto the HSP70 model revealed contacts involving both its nucleotide-binding domain (NBD) and substrate-binding domain (SBD) (Fig. 2F). On IFT172, HSP70 binding clustered at two discrete TPR subregions: Site 1 (∼K821-K913) and Site 2 (K1301-C-terminus). We validated this bipartite recognition mode by co-immunoprecipitation of IFT172 TPR fragments expressed in HEK293T cells, including the full TPR domain (residues 594-1749), Site 1 (residues 812-984), and Site 2 (residues 1275-1470). HSP70 bound robustly to constructs containing either Site 1 or Site 2, as well as the full-length TPR domain (Fig. 2G). Together, these data support a cooperative “spatial handoff” model (Fig. 2H), in which TRiC encapsulates the aggregation-prone WD40 domains within its folding chamber, while HSP70 simultaneously stabilizes the elongated TPR solenoid extending in the cytosol. This coordinated, spatially segregated chaperone action likely shields exposed hydrophobic surfaces from non-native interaction, ensuring efficient folding of the ∼200 kDa IFT172 polypeptide across two distinct chaperone environments.

### TRiC exhibits allosteric flexibility to accommodate oversized substrates

To understand how TRiC structurally accommodate such a massive substrate, we analyzed the conformational landscape of the apo-state TRiC-IFT172 complex by 3D variability analysis (3DVA) (Movie. S1). This revealed pronounced flexibility in the A-and I-domains of CCT2, CCT4, and CCT7, which undergo coordinated outward pivoting driven by coupled intra-ring (mode 1) and inter-ring (mode 2) dynamics, collectively expanding the central chamber. Focused 3D classification around the CCT7/2/4 subunits resolved these continuous motions into discrete conformational states characterized by pronounced bending of individual subunits (Fig. 3A, Fig. S2B, C). For example, the A- and I- domains of CCT4 bend outward by ∼81° relative to the equatorial base (Fig. 3C, CCT2 and CCT7 in Fig. S3B-C). This extreme Z-shaped architecture mirrors the CCT2 conformation observed in yeast TRiC in the nucleotide partially preloaded (NPP) state^30^ and resembles the CCT4 conformation recently observed in TRiC-HDAC3 complexes, where it is associated with prefoldin release^31^.

**Fig. 3.**
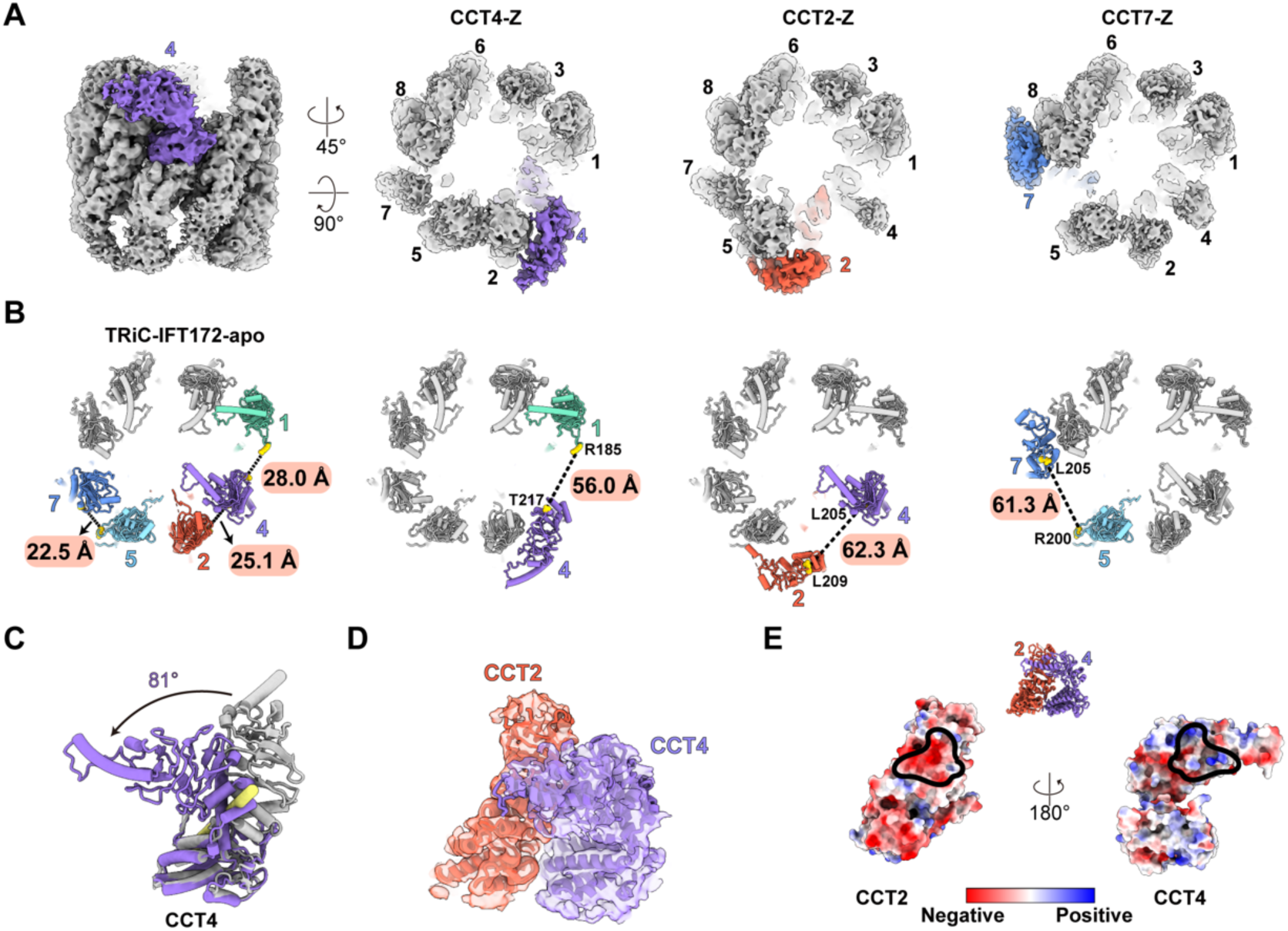
TRiC undergoes subunit-specific allosteric remodeling to accommodate an oversized substrate. (A) Cryo-EM classification of the TRiC-IFT172 complex resolves multiple conformational states in which the CCT2, CCT4, and CCT7 subunits adopt characteristic Z-shaped bending conformations. (B) Z-shaped remodeling expands the chamber aperture. Inter-subunit distances (CCT4-CCT1, CCT2-CCT4, and CCT7-CCT5) are shown before (left) and after bending, illustrating the magnitude of chamber widening. (C-E) Analysis of the CCT4-Z map: measurement of the CCT4 bending angle using an automatically defined rotation axis (in yellow) in ChimeraX (C); model-to-map fitting of the Z-shaped CCT4-CCT2 subunit pair (D); electrostatic surface potential analysis of the CCT4-CCT2 interface, suggesting stabilizing electrostatic complementarity in the Z-shape conformation (E). Black outlines depict their interaction surfaces.

This allosteric remodeling considerably widens the folding chamber aperture. The resulting “jaw-opening” movement expands inter-subunit distances by 28.0 Å (CCT4-CCT1), 37.2 Å (CCT2-CCT4), and 38.8 Å (CCT7-CCT5) (Fig. 3B). These expansions provide the steric clearance required for bulky substrates such as IFT172 to enter the chamber while allowing non-encapsulated regions, such as the TPR domain, to extend into the cytosol. That these bending events occur stochastically at different subunit positions suggests an adaptive mechanism in which TRiC local remodels its rim to match the footprint of individual clients. Structural analysis further indicates that these metastable bent conformations are stabilized by complementary electrostatic networks between the I-domains of the bent subunits and their neighboring “lean-on” partners (Fig. 3D-E, Fig. S3D-E).

Collectively, these findings reveal a fundamental structural plasticity in the eukaryotic chaperonin. Unlike the rigid, predetermined geometry of the GroEL-GroES cage, TRiC subunits possess an intrinsic capacity for transient, large-scale conformation rearrangements. This modular allostery enables the complex to break symmetry and dynamically resize its chamber, a feature that likely underlies TRiC’s ability to fold the diverse and structurally complex proteome of eukaryotic cells.

### Folding of IFT172 occurs in the open TRiC chamber, independent of ring closure

To visualize the folding trajectory of IFT172, we determined the cryo-EM structure of the TRiC-IFT172 complex in the presence of ATP-AlFx, a widely used ATP-hydrolysis transition-state analog that triggers TRiC ring closure^15,16,32,33^. The reconstruction resolved two states, an open state and a closed state (Fig. S5A-D). In the open state, well-resolved nucleotide density consistent with ATP was observed in all binding pockets (Fig. S5E). ATP binding stabilizes the complex relative to the apo state. 3DVA revealed that the CCT2, CCT4, and CCT7 retract from the pronounced outward-bending conformations observed in the apo-state (Movie. S2). Strikingly, a disk-shaped density was resolved above the disordered N- and C-terminal tails of TRiC, corresponding to a nearly folded WD40 domain within the open chamber (Fig. 4A-B).

**Fig. 4.**
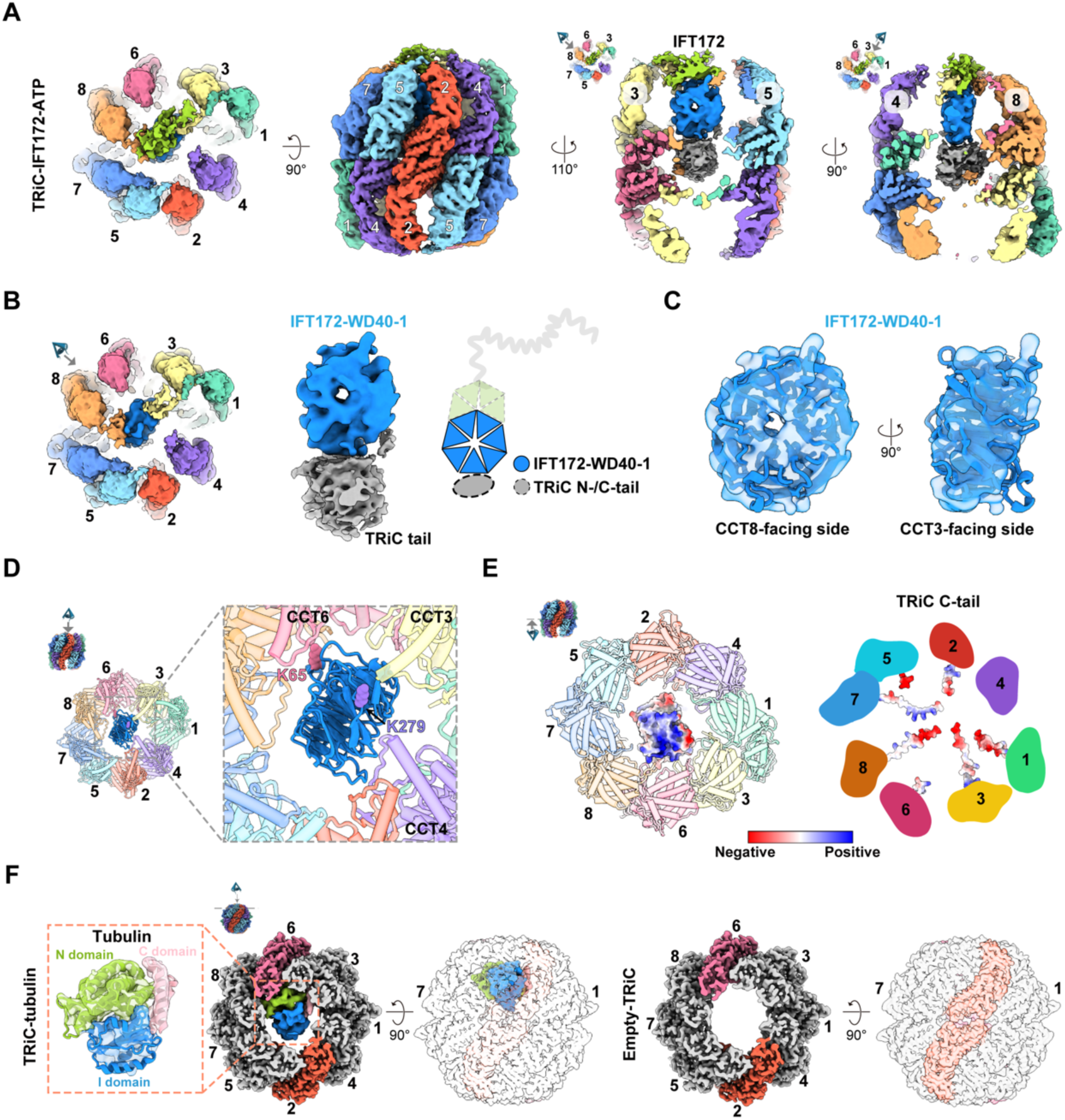
WD40-1 domain of IFT172 is folded in the open, ATP-bound TRiC chamber and is ejected upon ring closure. (A) Cryo-EM map of the ATP-bound TRiC-IFT172 complex, with the last two panels showing cut-away views of the center chamber with the blue density illustrating the WD40-1, yellow green density most likely present WD40-2, grey density corresponding to the disordered N-/C-terminal tails of TRiC. (B-C) Density corresponding to the folded WD40-1 domain of IFT172 (blue) within the open, ATP-bound TRiC chamber, positioned above the disordered N- and C-terminal tails of TRiC (grey), and interacting with the A domains of CCT3 and CCT8. (D) XL-MS restraints supporting the orientation of WD40-1 within the chamber, with K65 crosslinked to CCT6, and K279 crosslinked to CCT4. (E) Electrostatic surface potential analysis indicates that the predominantly negatively charged C-terminal tails of TRiC potentially engage the positively charged surface of IFT172 WD40-1 domain, stabilizing folding in the open state. (F) Cryo-EM analysis shows that in the closed TRiC chamber, IFT172 density is absent. The closed chamber is either occupied by tubulin (left three panels) or empty (right two panels). Tubulin domains are colored as indicated.

AlphaFold prediction indicates that, of the two IFT172 WD40 domains, only WD40-1 forms a canonical circular β-propeller. Consistent with this, XL-MS crosslinks under ATP-AlFx conditions mapped exclusively to the WD40-1 domain (Fig. S6A, Table. S4). We therefore assign the disk-shaped density to folded WD40-1. It is orientated with its top face toward CCT8, its bottom face toward CCT4, and its circumference positioned toward CCT3 and CCT5 (Fig. 4B-C). This orientation is supported by specific crosslinks, including K65 to CCT6 and K279 to CCT4 (Fig. 4D). The WD40-2 and TPR domains are not well-resolved in the map, consistent with a topology in which these regions extend outside the chamber.

Although hydrophobic shielding of inter-blade interfaces is a general requirement for WD40 β-propeller folding (Fig. S6B), our structure suggests that TRiC facilitates this process through electrostatic interactions. Specifically, WD40-1 rests primarily on a platform formed by the disordered N- and C-terminal tails of TRiC subunits (Fig. 4A, B), and electrostatic surface analysis indicates that its highly positively charged surface is complemented by the negatively charged, flexible C-termini of TRiC (Fig. 4E). This tentacle-like contact mode contrasts sharply with canonical substrates, including actin, tubulin, Gβ, HDAC3, and σ3, whose folding relies heavily on interactions with the rigid cavity walls^12,13,15–17,31^. Additional electrostatic stabilization is provided by the apical domains of CCT3 and CCT8 (Fig. S6C-D). Crucially, the observation of a folded WD40-1 domain within the open, ATP-bound chamber defines an unexpected folding intermediate and demonstrates that productive folding can occur prior to encapsulation, a phenomenon not observed with smaller TRiC clients.

Most strikingly, the closed TRiC chamber was completely absence of IFT172 density. Instead, closed chambers were either empty or occupied by a density corresponding to endogenous tubulin (Fig. S5F), as confirmed by high-resolution reconstruction of the cross-linked complex (Fig. 4F, Fig. S7). The presence of IFT172 in the open state but excluded from the closed state implies that ring closure acts as a mechanical trigger for substrate IFT172 ejection rather than encapsulation.

Together, these results define a non-canonical “fold-and-eject” trajectory for oversized substrates. Unlike small substrates (< 70 kDa), which require encapsulation within the closed TRiC chamber, IFT172 completes folding of its N-terminal WD40-1 domain in the open, ATP-bound state, where the flexible terminal tails and apical domains of TRiC collectively support maturation of the critical β-propeller. The subsequent transition to the closed state, driven by ATP hydrolysis, mechanically displaces the folded product and releases it for downstream assembly. This pathway resolves the steric paradox of how TRiC productively folds substrates that physically exceed the physical capacity of its closed chamber.

### TRiC is essential for functional IFT assembly *in vivo*

To assess the physiological relevance of this chaperonin pathway for intraflagellar transport (IFT), we established *C. elegans* as an *in vivo* model system. Using CRISPR/Cas9, we generated a strain expressing endogenous IFT172 (OSM-1) fused to GFP, enabling live-cell imaging of IFT dynamics in sensory cilia by confocal fluorescence microscopy^34^ (Fig. 5A). We then performed neuron-specific RNAi knockdown of the TRiC subunits *cct-5* and *cct-8* to evaluate the impact of chaperonin depletion on ciliary function.

**Fig. 5.**
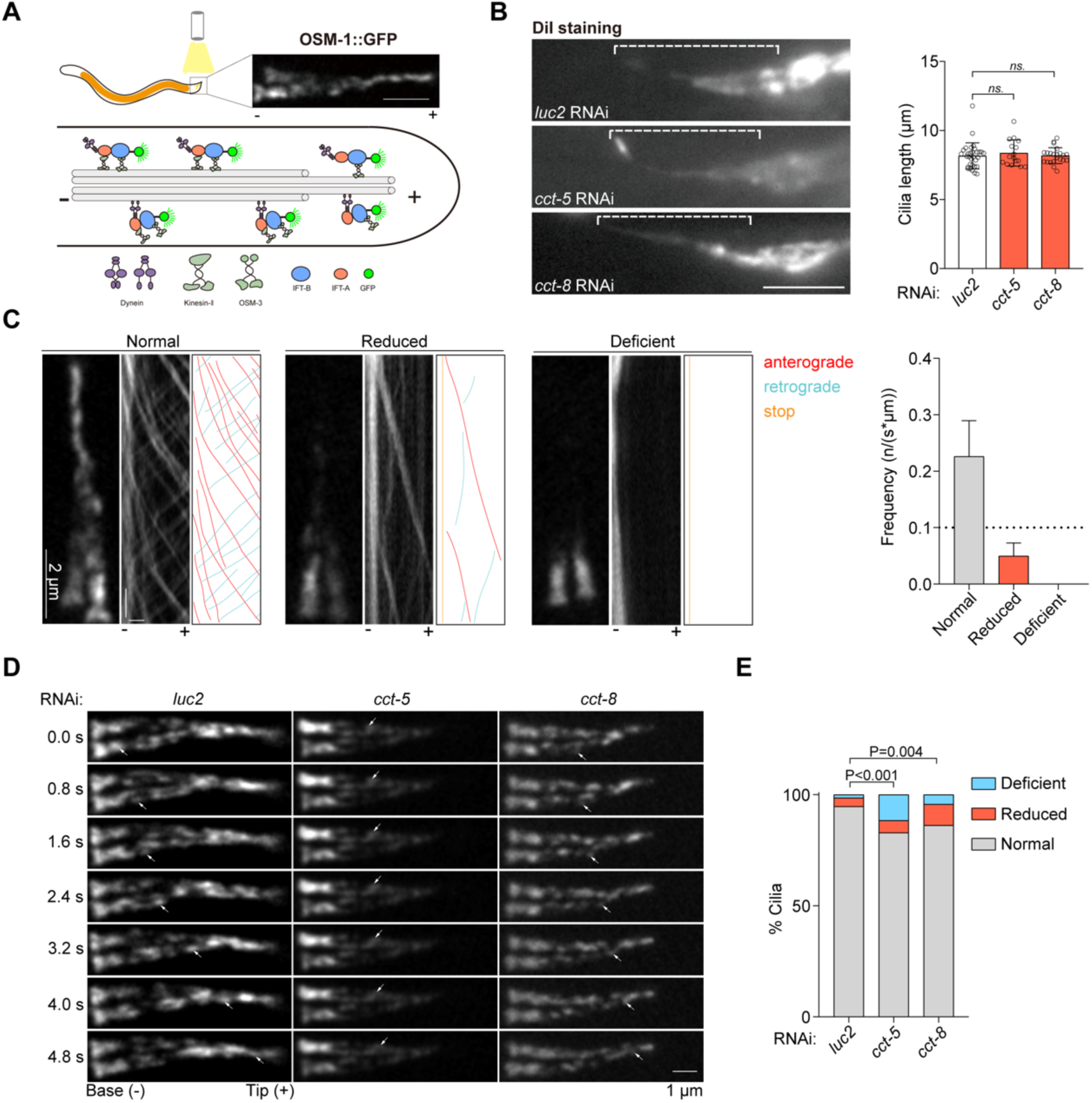
TRiC activity is required for efficient intraflagellar transport *in vivo*. (A) Schematic of intraflagellar transport (IFT) in sensory cilia located at the dendritic endings of *C. elegans* sensory neurons. “–” indicates the ciliary base, and “+” the ciliary tip. Endogenous IFT172 (OSM-1) was tagged with GFP by CRISPR/Cas9-mediated genome editing. (B) DiI staining of sensory neurons in worms subjected to neuron-specific RNAi against *luc2* (control, n = 31), *cct-5* (n = 16), or *cct-8* (n = 22). No significant differences in cilia length were observed among the groups. Scale bar, 2 μm. (C) Confocal live imaging of IFT dynamics in sensory cilia. RNAi-treatment of *cct-5* or *cct-8* resulted in a pronounced reduction or complete loss of IFT particle motility compared with control. Horizontal scale bar, 1 μm; vertical scale bar, 2 s. (D) Representative kymographs showing normal IFT trajectories in controls and diminished IFT events in RNAi treated worms. (E) Quantification of IFT defects in RNAi-treated worms using the Mann-Whitney U test. A total of 209 animals were analyzed for *luc2* RNAi, 164 for *cct-5* RNAi, and 189 for *cct-8* RNAi. Statistical significance is indicated by the P values shown above the histograms.

We first examined gross ciliary morphology using DiI staining, a lipophilic dye that labels the ciliary membrane. Neuron-specific knockdown of *cct-5* or *cct-8* did not significantly alter ciliary length compared with controls (Fig. 5B), suggesting that partial TRiC depletion does not abolish ciliogenesis. By contrast, quantitative analysis of IFT dynamics revealed a pronounced functional defect. In normal animals, OSM-1::GFP particles displayed robust bidirectional IFT transport along the axoneme. In TRiC-deficient neurons, however, transport activity was markedly reduced, and many cilia showed a complete loss of IFT events (Fig. 5C-D, Mov. S3). Quantification confirmed that knockdown of either *cct-5* or *cct-8* significantly increased the frequency of defective IFT phenotypes (Fig. 5E).

These data demonstrate that, although partial reduction of TRiC subunits may suffice to maintain axonemal structure, efficient and robust IFT requires full chaperonin capacity. This phenotype is consistent with our biochemical model: depletion of TRiC likely creates a bottleneck in the supply of folded IFT172, limiting its assembly into functional IFT trains and thereby staling transport without necessarily disassembling the existing axoneme.

### TRiC serves as a general folding hub for the IFT machinery

The tandem WD40-TPR architecture of IFT172 is not unique; it is a conserved structural motif shared by five other IFT subunits: IFT80 in the IFT-B complex, IFT121, IFT122, IFT140, and IFT144 in the IFT-A complex (Fig. 6A-B). This structural conservation raises the possibility that TRiC functions as a general folding hub for this class of large, multidomain ciliary proteins.

**Fig. 6.**
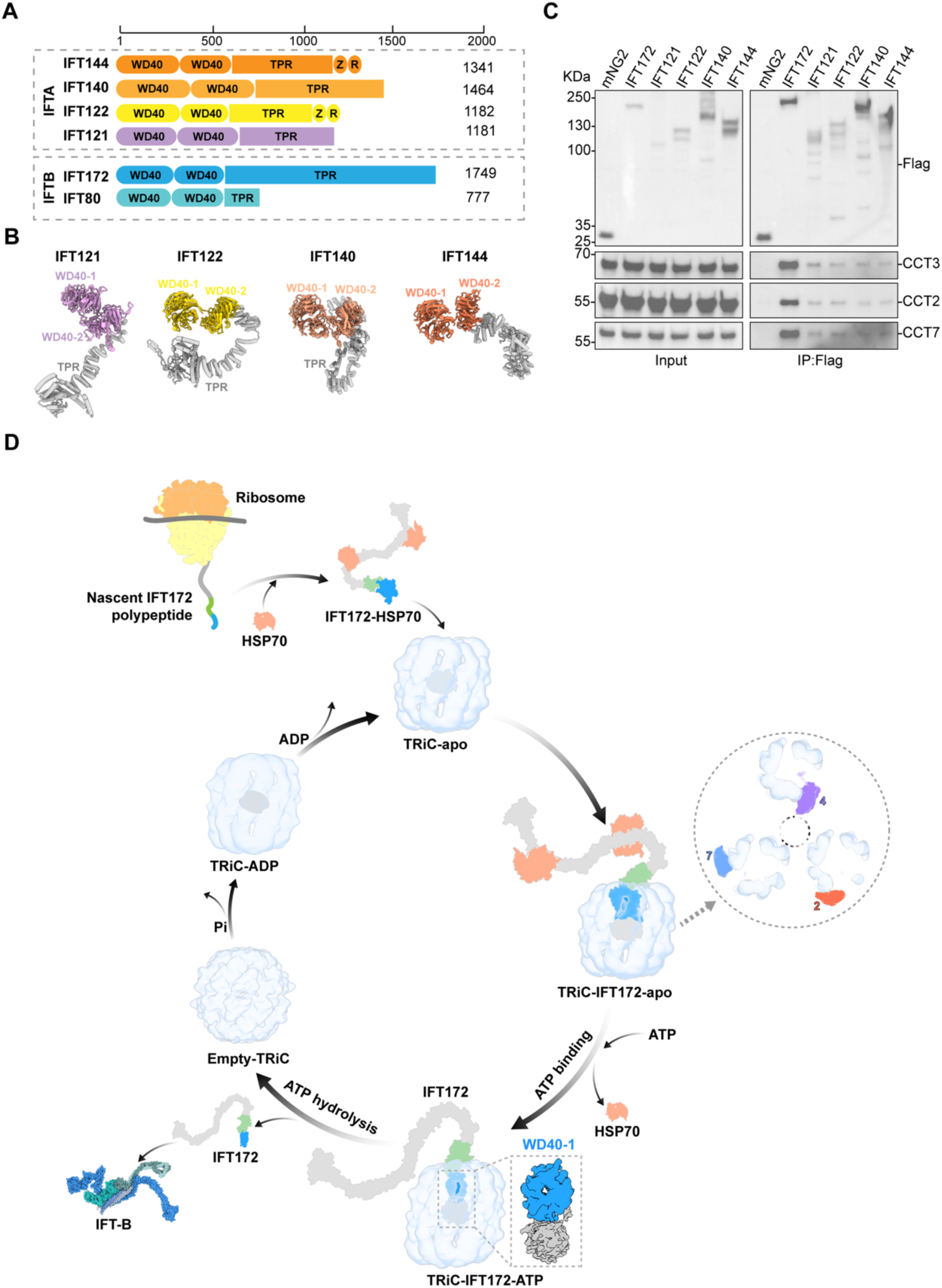
TRiC engages multiple IFT subunits with conserved tandem WD40-TPR architecture and folds oversized substrate via a non-canonical mechanism. (A) Domain organization of IFT-A and IFT-B subunits analyzed in this study. WD40, TPR, Z, and R denote WD40 β-propeller repeats, TPR-like domains, zinc-binding domains, and RING finger domains, respectively. (B) AlphaFold3-predicted structures of IFT121, IFT122, IFT140, and IFT144, highlighting the conserved tandem WD40 domains (colored) and C-terminal TPR-like domains (gray). (C) Co-immunoprecipitation of transiently expressed FLAG-tagged IFT subunits (IFT172, IFT121, IFT122, IFT140, and IFT144) from HEK293T cells shows interactions with TRiC subunits (CCT2, CCT3, and CCT7). (n = 3). (D) Working model for TRiC-mediated folding of IFT172 and other oversized, multi-domain IFT substrates. Upon release from the ribosome, the nascent IFT172 polypeptide is bound by HSP70. The apo-state TRiC then sequesters the aggregation-prone WD40 domain within its central chamber, while HSP70 continues to stabilize the exposed distal segments in the cytosol. Notably, TRiC subunits CCT2/4/7 adopt an extended Z-shaped conformation, expanding the chamber to accommodate the bulky multi-domain substrate. Subsequent ATP binding promotes HSP70 dissociation and drives an allosteric rearrangement of TRiC, facilitating the folding of the IFT172 WD40-1 domain in an open chamber. Following ATP hydrolysis, chamber closure triggers the release of folded IFT172, permitting its incorporation into the IFT-B complex via a “fold-and-eject” mechanism. TRiC subsequently releases Pi and ADP, reverting to the apo-state to initiate a new cycle. The TRiC-ADP model is based on the human TRiC-ADP map (EMD-32993).

To test this hypothesis, we performed co-immunoprecipitation with FLAG-tagged IFT-A subunits (IFT121, IFT122, IFT140, and IFT144). Remarkably, all tested IFT-A components co-purified with endogenous TRiC subunits (CCT2, CCT3, CCT7) (Fig. 6C). Although these interactions appeared weaker than that of IFT172, suggesting that IFT172 may be the most kinetically demanding or aggregation-prone client in this family, the conserved TRiC association across both IFT-A and IFT-B subunits points to a shared dependence on the chaperonin for biogenesis. Collectively, these results establish TRiC as a central biogenesis hub for the IFT machinery. By recognizing the conserved WD40 β-propeller topology, TRiC likely orchestrates the folding of multiple core components across both IFT-A and IFT-B complexes, thereby safeguarding the structural and functional integrity of the ciliary transport system.

## Discussion

This study addresses a longstanding paradox in chaperone biology: how the eukaryotic chaperonin TRiC folds substrates that exceed the capacity of its closed chamber. By delineating the folding pathway of IFT172, the largest subunit (∼200 kDa) of the intraflagellar transport machinery, we uncover a “divide-and-conquer” strategy in which TRiC and HSP70 simultaneously engage distinct structural domains of the same client (Fig. 6D). Structurally, TRiC undergoes pronounced Z-shaped bending to accommodate the WD40 domains, after which WD40-1 domain reaches a near-folded state within the open, ATP-bound TRiC chamber. Subsequent TRiC ring closure triggers substrate ejection rather than encapsulation, establishing a non-canonical “fold-and-eject” mechanism that resolves the size paradox posed by large TRiC substrates. We further show that this pathway is essential for ciliary function *in vivo* and conserved across multiple IFT subunits sharing tandem WD40-TPR architecture. Together, these findings establish a new paradigm for chaperonin-assisted folding of oversized multi-domain proteins, and reveal TRiC as a central proteostasis hub for ciliary protein biogenesis, with direct implications for ciliopathy pathogenesis.

The folding of multi-domain proteins that exceed the volumetric capacity of a single chaperone poses a fundamental challenge for the cell. For IFT172, we show that this challenge is met through a coordinated “spatial handoff”: TRiC captures the aggregation-prone WD40 domains in its chamber, whereas HSP70 independently holds and stabilizes the extending TPR solenoid in the cytosol (Fig. 2H). This modular cooperation effectively decouples the folding of individual domains from the steric constraints imposed by the full-length polypeptide, solving the volume problem by ensuring that only the most aggregation-prone regions require encapsulation by TRiC.

In contrast to the co-translational models in which chaperones and chaperonins act sequentially on nascent chains^35,36^, our findings reveal a post-translational mode of cooperation in which the two systems operate in parallel on distinct domains of the same polypeptide. This “divide-and-conquer” strategy may therefore represent a general principle for the biogenesis of large, multi-domain eukaryotic proteins, extending the functional range of the chaperonin system beyond its physical size limits.

An interesting implication of this work is that TRiC exploits intrinsic structural plasticity to accommodate oversized substrates. Our cryo-EM analysis reveals that specific subunits (CCT2, CCT4, CCT7) undergo dramatic Z-shaped outward bending of the A-/I-domains, up to ∼97° for CCT7, widening the chamber aperture by up to 38.8 Å (Fig. 3). These conformations resemble the CCT2 Z-shape observed in yeast TRiC^30^ and the CCT4 bending seen in TRiC–HDAC3 complexes^31^, suggesting that this plasticity reflects a conserved feature of the chaperonin rather than a substrate-specific exception. The stochastic occurrence of bending at different subunit positions further suggests an adaptive mechanism in which TRiC locally remodels its rim to match the footprint of individual clients. This symmetry-breaking capacity stands in sharp contrast to the rigid geometry of the prokaryotic GroEL-GroES system and likely represents a key evolutionary adaptation for servicing the structurally diverse eukaryotic proteome.

Our most unexpected findings is that IFT172’s WD40-1 domain achieves a near-native fold within the open, ATP-bound TRiC chamber, before ring closure, challenging the canonical “folding-upon-closure” dogma^18^ in which encapsulation within the closed chaperonin chamber is required for productive folding^12–17^. Critically, this folding is mediated by interactions with TRiC’s flexible terminal tails and apical domains (Fig. 4A-E), a tentacle-like contact mode distinct from the rigid cavity-wall interactions that govern folding of smaller clients such as actin, tubulin, and Gβ^12–17^. Subsequent ring closure ejects the nearly folded product rather than encapsulating it (Fig. 4F), assigning distinct roles to the two ATPase states: the open ATP-bound state serves as the folding environment, whereas closure acts as a mechanical release trigger. This “fold-and-eject” mechanism resolves the steric paradox of how TRiC folds substates that exceed the volume of its closed cavity and establishes that chaperonin folding trajectories are substrate-dependent rather than uniform, a principle that may extend to other large, multi-domain TRiC clients.

Beyond advancing our understanding of chaperonin mechanism, this work has direct implications for ciliary biology and ciliopathy pathogenesis. The tandem WD40-TPR architecture of IFT172 is conserved across five additional IFT subunits spanning both IFT-A and IFT-B subcomplexes (Fig. 6A-B). We show that TRiC engagement is a shared requirement for this protein family (Fig. 6C), positioning TRiC as a central biogenesis hub for the IFT machinery. The ciliopathy-associated IFT172 I411N mutation impairs TRiC binding (Fig. 2E), directly linking defective chaperonin recognition to human disease and suggesting that some ciliopathy mutations may act by compromising chaperone engagement rather than native protein function per se. Furthermore, neuron-specific partial TRiC depletion in *C. elegans* greatly reduces IFT transport activity without disrupting axonemal structure (Fig. 5), demonstrating that TRiC is essential for functional IFT assembly *in vivo*. The conserved TRiC dependence across multiple IFT components raises the possibility that perturbations in chaperonin capacity, whether caused by mutation, aging, or proteotoxic stress, could simultaneously compromise the entire IFT system.

In summary, we delineate a complete TRiC-assisted folding pathway for the giant ciliary protein IFT172, and uncover three mechanistic principles that establish a new framework for handling oversized substrates: domain-partitioned chaperone cooperation, dynamic chamber remodeling, and ATP-bound open-state folding coupled to closure-triggered ejection. These findings establish a new framework for how chaperonin networks handle oversized substrates, expand the known repertoire of chaperonin folding mechanisms, and identify TRiC as a critical proteostasis node for ciliary biogenesis, with implications for understanding and potentially treating ciliopathies driven by defective protein folding.

## Methods and materials

### Plasmids construction

Full-length mouse *Ift172* (NM_026298.5) was amplified from mouse cDNA and cloned into the pcDNA3.4 vector with a C-terminal FLAG tag via homologous recombination. Truncation constructs, including WD40-1 (residues 1–316), WD40-2 (317–593), TPR (594–1749), TPR-Site1 (814–984), and TPR-Site2 (1275–1470) were cloned into pcDNA3.4 with an N-terminal FLAG tag followed by a TEV cleavage site. Disease-associated point mutants were generated via site-directed mutagenesis on the C-terminally FLAG-tagged full-length template. Coding sequences for mouse IFT-A subunits (*IFT121*: NM_172470.3; *IFT122*: NM_031177.4; *IFT140*: NM_001410319.1; *IFT144*: NM_153391.2) were obtained from Youbao Biotech and cloned into pcDNA3.4 via homologous recombination with a C-terminal FLAG tag.

### Cell culture and transfection

HEK293T cells were maintained in DMEM supplemented with 10% (v/v) fetal bovine serum (FBS) and 1% (v/v) penicillin-streptomycin at 37 °C under 5% CO_2_. For transient transfection, cells were seeded in 6-well plates and cultured for 18–24 h to reach ∼80% confluency. Plasmid DNA was transfected using Lipo8000 (Beyotime) according to the manufacturer’s instructions. Cells were harvested 48 h post-transfection and processed immediately for immunoprecipitation.

Expi293F cells were cultured in serum-free 293F medium at 37 °C in a humidified shaking incubator (130 rpm, 5% CO_2_). Transient transfections were performed using polyethyleneimine (PEI), with a DNA:PEI ratio of 1:3 (w/w). Cells were collected 48 h post-transfection by centrifugation, washed once with PBS, and resuspended for subsequent protein purification.

### Purification of the TRiC-IFT172 complex

Expi293F cells overexpressing FLAG-tagged IFT172 were harvested by centrifugation and resuspended in lysis buffer (20 mM HEPES pH 7.5, 10 mM KCl, 1.5 mM MgCl_2_, 250 mM sucrose, 5 mM β-ME, 0.15% DDM), supplemented with one tablet of protease inhibitor cocktail tablet (Roche) per 100 mL of lysate. Cell lysis was carried out for 30 min at 4 °C with gentle agitation. The lysate was clarified by centrifugation at 750 × g for 10 min to remove unbroken cells and nuclei. NaCl was then added to the supernatant to a final concentration of 150 mM, followed by a second centrifugation at 18,000 rpm for 30 min at 4 °C to remove insoluble debris.

The clarified supernatant was incubated with 1 mL of anti-FLAG affinity resin (Anti-DYKDDDK Tag Affinity Gel; Smart-lifesciences) for 3 h at 4 °C with gentle rotation. The resin-lysate mixture was loaded onto a gravity-flow column and washed with 50 column volumes of wash buffer (20 mM HEPES pH 7.5, 150 mM NaCl, 5 mM β-ME, 10% glycereol, 0.15% DDM). Bound proteins were eluted using the same buffer supplemented with 0.2 mg/mL 3×FLAG peptide (Sigma). Eluted fractions were concentrated using a 100 kDa molecular weight cutoff centrifugal filter (Amicon Ultra, Millipore) and further separated by 10-40% (v/v) glycerol gradient centrifugation (20 mM HEPES pH 7.5, 150 mM NaCl, 1 mM DTT). Fractions containing the TRiC-IFT172 complex were identified, combined, and concentrated using a 100 kDa cut-off centrifugal filter. The sample was subsequently exchanged through multiple rounds of concentration and dilution into MQA buffer (50 mM NaCl, 20 mM HEPES pH 7.4, 5 mM MgCl_2_, 0.1 mM EDTA, 1 mM DTT, 5% glycerol) for downstream biochemical and structural analyses.

### Cryo-EM grids preparation

To prepare the TRiC-IFT172 vitrified sample, the purified TRiC-IFT172 complex was diluted to 1.5 mg/mL, and crosslinked with 0.05% glutaraldehyde for 1 h on ice. The reaction was quenched by addition of 100 mM Tris-HCl for 10 min. An aliquot of 2 μl of sample was applied to poly-Lysine pretreated holey carbon grid (Quantifoil R1.2/1.3 or R2/1, Cu, 200 mesh), blotted using a Vitrobot Mark IV (Thermo Fisher Scientific), and plunge-frozen in liquid ethane cooled by liquid nitrogen. For ATP-AlFx condition, purified TRiC-IFT172 was incubated with 5 mM MgCl_2_, 5 mM Al (NO_3_)_3_, 30 mM NaF, and 1 mM ATP at 37 °C for 1 h to generate the ATP-hydrolysis transition-state. To further stabilize the transition-state complex, a crosslinked sample was treated with 0.1% glutaraldehyde for 1 h, quenched with 100 mM Tris-HCl, and vitrified as described above.

### Cryo-EM data acquisition

Cryo-EM movies of the TRiC-IFT172 sample were collected on a Titan Krios (Thermo Fisher Scientific) operated at an accelerating voltage of 300 kV. Data were acquired in automated mode using EPU software (Thermo Fisher Scientific) on a K3 direct electron detector (Gatan) in counting mode at 64,000× nominal magnification (yielding a pixel size of 1.093 Å). Each frame was exposed for 0.1 s, and the total accumulation time was 3 s, corresponding to a total dose of 50.2 e^−^/Å^2^ on the specimen.

For the TRiC-IFT172-ATP-AlFx sample, movies were collected on a Titan Krios using a K3 detector in counting mode (pixel size: 0.86 Å) with EPU automation software. Each frame was exposed for 0.1 s, and the total accumulation time was 3 s, leading to a total dose of 56.8 e^−^/Å^2^ on the specimen.

For the crosslinked TRiC-IFT172-ATP-AlFx sample, movies were acquired on a Glacios2 electron microscope (Thermo Fisher Scientific) operated at 200 kV using a Falcon 4i detector (Thermo fisher Scientific) in counting mode (yielding a pixel size of 0.91 Å) using EPU software. The total accumulated dose was 50.0 e^−^/Å^2^ in total accumulation time of 4.42 s on the specimen.

### Image processing and reconstruction

Single-particle analysis was performed using RELION 4.0^37^ and cryoSPARC 4.2.1^38^. Movie frames were aligned and summed using MotionCorr2 whole-image motion correction software^39^. CTF parameters were estimated using CTFFIND4^40^, and particles were autopicked using crYOLO 1.7.6^41^.

For the TRiC-IFT172 dataset, 1,655,238 particles were picked from 8,822 micrographs. After 2D classification, 1,040,775 particles were retained and subjected to heterogeneous refinement, yielding 435,781 particles displaying good features of open-state TRiC. Following CTF refinement, Bayesian polishing, and non-uniform refinement, a consensus map at 3.41 Å resolution was obtained. To improve the substrate density, these 435,781 particles were subjected to focused classification on the chamber. Particles exhibiting substrate density in the trans ring were rotated by 180° about the Z-axis to align with those containing substrate in the cis ring, thereby resolving the cis/trans ring ambiguity and yielding 372,833 particles. A second round of focused classification on the substrate region produced 219,132 particles, which were locally refined to generated a 3.55 Å resolution map of the TRiC-IFT172 complex. The final map was further processed using normfilt function in EMAN1^42^ to enhance substrate continuity. Further 3DVA of this dataset indicated conformational flexibility primarily in CCT7, CCT2, and CCT4 subunits. To resolve these conformational states, the 435,781 apo-state particles were refined with C2 symmetry, followed by symmetry expansion to align both half-rings to one side, effectively doubling the particle number. Multiple rounds of focused no-alignment 3D classification then yielded reconstructions of CCT7-Z (57,802 particles; 3.81 Å resolution), CCT2-Z (38,426 particles, 3.89 Å), and CCT4-Z (58,082 particles, 3.82 Å) states.

For the TRiC-IFT172-ATP-AlFx dataset, from 23,458 micrographs, 4,508,286 particles were selected after 2D classification. 3D classification separated 1,443,227 open-state particles and 1,032,616 closed-state particles, yielding consensus maps at 3.09 Å and 2.83 Å resolution, respectively. To resolve extra density in the open chamber, focused 3D classification was performed using a chamber mask; cis/trans ambiguity was corrected by rotating trans-ring substrate particles by 180° about the Z-axis, yielding 573,318 particles with discernible IFT172 density. Focused classification on the cis-ring substrate region produced 105,469 particles displaying clear WD40 features, which were refined after CTF refinement and Bayesian polishing to a 3.61 Å resolution map. For the 1,032,616 closed-state particles, focused 3D classification using a mask on the CCT3/6/8 A-domains separated particles by cis-ring occupancy. Two additional rounds of tandem focused classification on the entire trans- and cis-ring chambers yielded 207,424 particles with extra density, refined to a 3.03 Å resolution map.

For the TRiC-IFT172-ATP-AlFx crosslinked dataset, from 66,373 micrographs, 3,087,621 particles were selected after 2D classification and subjected to heterogeneous refinement, yielding 823,779 open-state and 1,754,599 closed-state particles. Closed-state particles were subjected to focused no-alignment 3D classification using a mask on the cis-ring CCT3/6/8 chamber region, separating cis-ring well-occupied (739,833 particles), partially occupied (298,056 particles), and empty (716,709 particles) classes. Chamber compositions were further assessed by focused classification using a trans-ring mask, revealing that ∼24% of closed-state particles were empty, while the remainder contained substrate density in various folding states. Focused classification on the full cis-ring resolved various tubulin folding states, rather than IFT172. The TRiC-tubulin and empty-TRiC complexes were refined individually to 4.37 Å and 4.40 Å, respectively, maps were sharpened using EMready^43^.

### Atomic model building

Atomic models of human TRiC in different open conformational states were built using the human TRiC structure (PDB: 7X0A) as the initial template^16^. Z-shaped conformations CCT2, CCT4, and CCT7 were generated by homology modelling in SWISS-MODEL using the yeast CCT2 Z-shaped conformation (PDB: 5GW4) as a reference^30^. Closed state TRiC model was generated by using human TRiC-ADP-AlFx structure (PDB: 7X0V) as template^16^. The IFT172 model was obtained from Alphafold3 (mostly using the *Chlamydomonas reinhardtii* IFT172 as template). Models were docked into cryo-EM maps via rigid-body fitting in UCSF Chimera^44^, followed by flexible fitting using Rosetta^45^. Iterative refinement was performed by manual adjustment in COOT and real-space refinement in Phenix (phenix.real_space_refine)^46^. Model geometry and fit-to-map statistics were validated in Phenix. Figures and electrostatic surface potential analyses were generated using UCSF Chimera and ChimeraX^47^.

### Cross linking mass spectrometry (XL-MS)

Purified complexes were crosslinked with BS^3^ (Sigma; spacer arm of 11.4 Å) at a final concentration of 4 mM on ice for 2 h. Reactions were quenched with 50 mM Tris-HCl (pH 7.5) for 15 min at room temperature. For ATP-AlFx condition, purified TRiC-IFT172 was incubated with 1 mM ATP, 5 mM MgCl_2_, 5 mM Al(NO_3_)_3_, and 30 mM NaF at 37 °C for 1 h prior to BS^3^ crosslinking as above. Crosslinked complexes were precipitated, resuspended, and digested with trypsin (enzyme-to-substrate ratio of 1:50 w/w) for 16 h at 37 °C. The tryptic-digested peptides were desalted and separated on an in-house packed capillary reverse-phase C18 column (40 cm length; 100 μm ID × 360 μm OD; 1.9-μm particle size; 120 Å pore diameter) coupled to an Easy LC 1200 system (Thermo Fisher Scientific). A 120 min-HPLC gradient of 6-35% buffer B (buffer A: 0.1% formic acid in water; buffer B: 0.1% formic acid in 80% acetonitrile) was applied at 300 nL/min. Eluting peptides were ionized and directly introduced into a Q-Exactive mass spectrometer (Thermo Fisher Scientific) equipped with a nano-spray source. Survey full-scan MS spectra (from m/z 300 to 1800) were acquired using an Orbitrap analyzer with a resolution r = 70,000 at an m/z of 400. Cross-linked peptides were identified and evaluated using pLink2 software^48^. Inter-protein crosslinks were filtered to require ≥ 2 observed spectra and an E-value ≤ 0.01.

### Structure prediction integrating cross-linking mass spectrometry data

To model the IFT172-HSP70 interaction in the apo state, we integrated cross-linking mass spectrometry (XL-MS) restraints with AlphaFold3 structure predictions. To calibrate distance constraints, we first analyzed Cα-Cα distances between cross-linked lysine pairs within the TRiC complex using its known structure (PDB: 7X0A), finding that 92% of lysine-lysine (K-K) pairs fell within 35 Å. We therefore adopted 35 Å as the maximum permissible distance for a valid cross-link restraint.

A total of 13 intermolecular K-K pairs were identified between HSP70 and IFT172. Predicted models were filtered to retain only those satisfying at least two cross-link restraints (< 35 Å). To cluster the remaining models, we constructed a binary satisfaction vector for each predicted structure, assigning a value of 1 if a specific K-K pair distance was < 35 Å and 0 otherwise. Models were then grouped into clusters based on a cosine similarity threshold of >0.9 between their vectors, yielding 10 clusters. We focused on the most highly populated clusters and manually screened the top 10 models from each cluster. This analysis identified two distinct HSP70-binding sites on the TPR domain of IFT172.

### Immunoprecipitation and Western blot

For co-immunoprecipitation, FLAG-tagged IFT172 construct (full-length, truncations, and point mutants) or FLAG-tagged IFT-A subunits were transiently expressed in HEK293T cells for 48 h. Cells were washed with PBS and lysed for 30 min in lysis buffer (25 mM HEPES pH 7.5, 150 mM NaCl, 2 mM MgCl_2_, 1 mM EGTA, 1 mM DTT, 10% glycerol, 0.5% NP40) supplemented with protease inhibitor cocktail. After lysate clearing by centrifugation, clarified protein extract was incubated with 20 μl packed anti-FLAG beads and incubated for 2 h at 4 °C. Beads were washed seven times with wash buffer (25 mM HEPES pH 7.5, 300 mM NaCl, 2 mM MgCl_2_, 1 mM EGTA, 1 mM DTT, 10% glycerol, 0.1% NP40) and boiled in SDS sample buffer.

For western blotting, input and IP samples were loaded onto 4–20% gradient gels (MOPS running buffer; ACE) and transferred to PVDF membranes (Millipore). Membranes were probed with the following primary antibodies: rabbit-FLAG (Sigma, F7425; 1:2000); rabbit-CCT2 (Abcam, ab92746; 1:5000); rabbit-GAPDH (PTM-5375, 1:5000); rabbit-HSPA1B (ABclonal, A19317, 1:1000).

### NADH-coupled ATPase assay of TRiC

TRiC ATP-hydrolysis rate was quantified using an NADH-coupled enzymatic assay^49^, in which each ATP-hydrolysis event allows for a pyruvate kinase (PK)-catalyzed conversion of one molecule of phosphoenolpyruvate into pyruvate, with pyruvate then converted to lactate by L-lactate dehydrogenase, resulting in oxidation of a single NADH molecule. Reactions were performed at 37°C in assay buffer (10 mM HEPES pH 7.5, 50 mM NaCl, 5 mM MgCl_2_, and 5% glycerol, 1 mM ATP). Experiments were performed in triplicate using 0.2 μM TRiC-IFT172 (or TRiC complex, as indicated). NADH consumption was monitored by measuring absorbance at 340 nm in 200 μl reaction volume in a 96-well plate reader. Data analysis was performed using GraphPad Prism 10.

### *C. elegans* RNAi, ciliary staining, and live imaging

*C. elegans*. RNAi experiments were performed by bacterial feeding on standard NGM plates containing IPTG using *E. coli* HT115, as described previously^50^. Worms were grown from the egg stage to day 1 adult on HT115 expressing dsRNA against the indicated genes. HT115 *[L4440::luc2]* was used as a negative control to minimize off-target effects, as *luc2* is not encoded in the *C. elegans* genome. The *[L4440::luc2]* strain was provided by the Antebi lab (MPI-AGE). The *[L4440::cct-5]* strain was a gift from the Jingdong Han lab (Peking University), and the *[L4440::cct-8]* RNAi construct was constructed using standard cloning methods, the primers used were: forward, 5′-ATGGCTATGAAGATCCCAAAG-3′; reverse, 5′-TTAGGCCATTCCATCGTCATC-3′. For neuron-specific RNAi, the *osm-1(syb7439[osm-1::GSGSG::gfp]) X.* strain (a gift from the Guangshuo Ou lab, Tsinghua University) was crossed with TU3401 (*sid-1(pk3321) V; uIs69 V.*) to enable tissue-restricted RNAi. The TU3401 strain was obtained from the Caenorhabditis Genetics Center (CGC).

DiI staining was performed as described^51^ with minor modifications. Day 1 adults were incubated with 1.7 μg/mL fluorescent Dye (DiI 1,1′-dioctadecyl-3,3,3′,3′,-tetramethylindo-carbocyanine perchlorate, Sigma) for 30 min at room temperature in dark, subsequently washed with M9 buffer, transferred to regular NGM plates for 30 min or more, and imaged on an Olympus BX53 microscope.

Live IFT imaging was performed as previous described^52^. Briefly, day 1 adults were anaesthetised with 16.7 mM levamisole in M9 buffer, mounted on 5% agar pads, and kept at room temperature. Phasmid cilia were imaged on an Olympus spinSR microscope with SoRa mode. Time-lapse images were acquired at an exposure time of 200 ms for 20s. Kymographs were generated and analyzed as described^52^. In brief, Fourier filtered and separated into anterograde and retrograde kymographs were generated with the KymographClear toolset plugin in ImageJ (http://www.nat.vu.nl/~erwinp/downloads.html).

### Statistics and Reproducibility

Cryo-EM data collection and processing statistics are summarized in Table. S1. Statistical analysis and graphing were performed in GraphPad Prism 10. Immunoblot band intensities were quantified using Image J. All biochemical assays were performed as three independent experiments.

Quantitative data are presented as mean ± standard deviation. Statistical significance is indicated as follows: ns (not significant), p > 0.05; *, p < 0.05; **, p < 0.01; ***, p < 0.001; ****, p < 0.0001.

All data presented in this study are available within figures and in Supplementary Information. Cryo-EM maps and atomic models have been deposited in the Electron Microscopy Data Bank (EMDB) and Protein Data Bank (PDB) under accession codes EMDB-XXXX/PDB-XXXX (to be provided) for each reconstruction. Detailed information is referred to the Table S1.

## Acknowledgements

We thank the staff members at the National Facility for Protein Science in Shanghai (https://cstr.cn/31129.02.NFPS) Electron Microscopy System, Mass Spectrometry System, Database and Computing System, and Protein Expression and Purification System for their instrument support and technical assistance. We are also grateful to Thermo Fisher Scientific for their support in cryo-EM data collection. This work was supported by grants from the Key Program of NSFC (32130056), National Key R&D Program of China (2024YFA1306204 and 2024YFA1803102), Strategic Priority Research Program of CAS (XDB0570303), general Program of NSFC (32570924), and Shanghai Pilot Program for Basic Research from CAS (JCYJ-SHFY-2022-008).

## Author contributions

Q.Z. and Y.C. conceived the study and designed the project. Q.Z. performed biochemical experiments, purified proteins, and conducted cryo-EM data collection and processing. J.L. performed *C. elegans* TRiC knockdown experiments and IFT imaging under the supervision of Y.S. Y.T. performed biochemical assays. Y. Y. helped with XL-MS analysis. Y.L. performed structural modeling based on XL-MS data under the guidance of S.H. W.H., Z.L., and Y.T. assisted with cryo-EM data collection. Y.W., J.F., W.J., and Q.S. assisted with biochemical experiments. Q.Z. and Y.C. wrote the manuscript with input from J.L., Y.S.

---

**Fig. S1.**
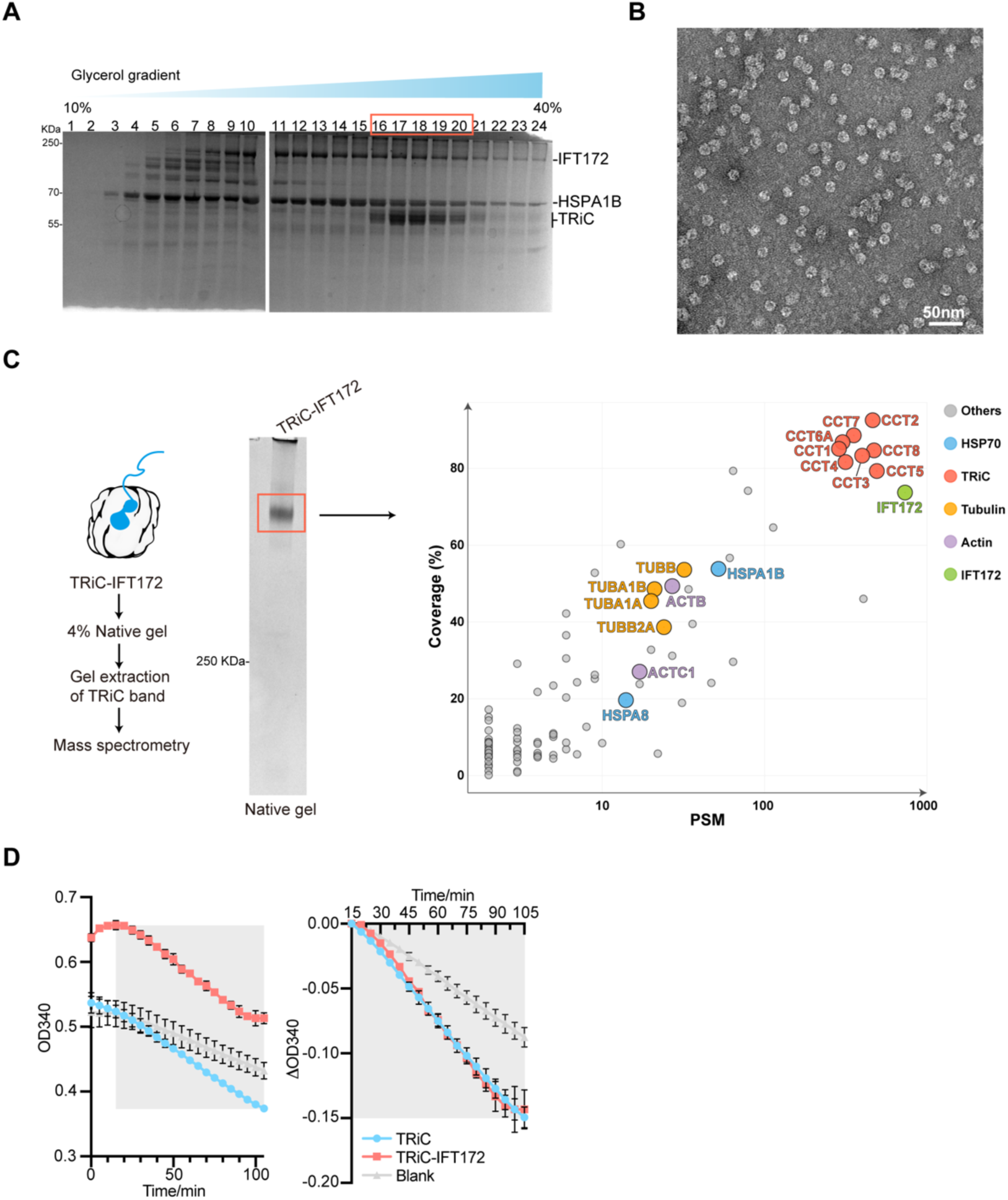
Purification and initial characterization of the TRiC-IFT172 complex. (A) Coomassie blue-stained SDS-PAGE of FLAG-IFT172 affinity purification followed by 10%–40% glycerol gradient centrifugation. Fractions 16–20 were pooled as the TRiC-IFT172 complex; bands corresponding to IFT172, TRiC subunits, and HSPA1B are indicated. (B) Representative negative-stain electron microscopy (NS-EM) micrograph of the purified TRiC-IFT172 complex. (C) Compositional analysis of the purified complex by mass spectrometry. The sample was resolved by 4% native PAGE; the TRiC-containing band was excised and subjected to mass spectrometry to identify co-purified proteins. (D) NADH-coupled ATPase assay showing that the ATP hydrolysis rate of the TRiC-IFT172 complex is comparable to that of TRiC alone.

**Fig. S2.**
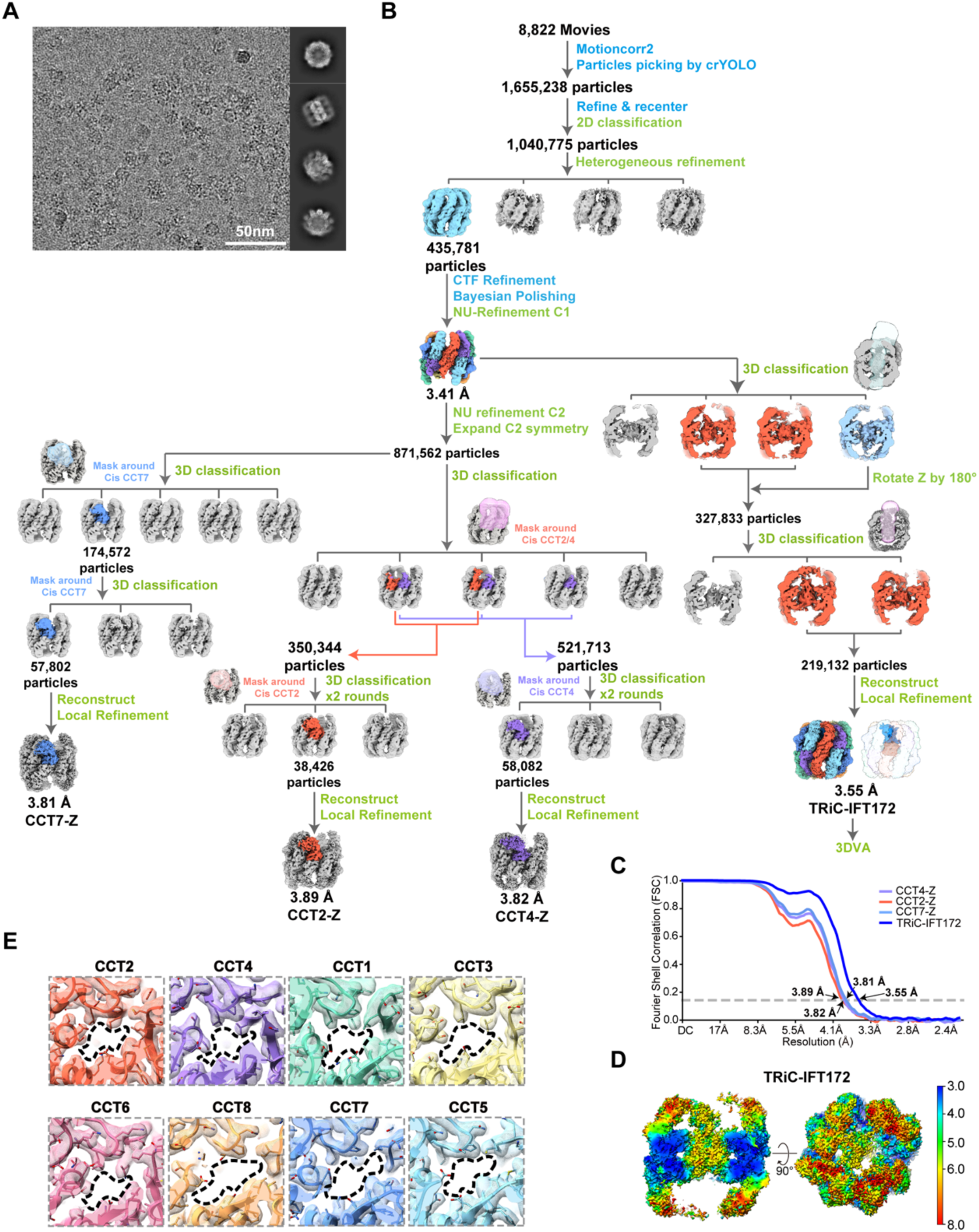
Cryo-EM data process and validation for the TRiC-IFT172 complex. (A) Representative cryo-EM micrograph and reference-free 2D class averages of the TRiC-IFT172 complex. (B) Cryo-EM single-particle processing workflow for the TRiC-IFT172 complex. Steps performed in RELION are indicated in blue and those in CryoSPARC in green. This color scheme is used consistently in subsequent workflow figures. (C) Fourier shell correlation (FSC) curves for the TRiC-IFT172 map and the CCT2-Z, CCT4-Z, and CCT7-Z conformational states. Resolutions were determined using the FSC = 0.143 criterion. (D) Local resolution estimations for the TRiC-IFT172 reconstruction. (E) Nucleotide-binding pockets analysis indicating that TRiC is in the apo state without bound nucleotide in the pockets.

**Fig. S3.**
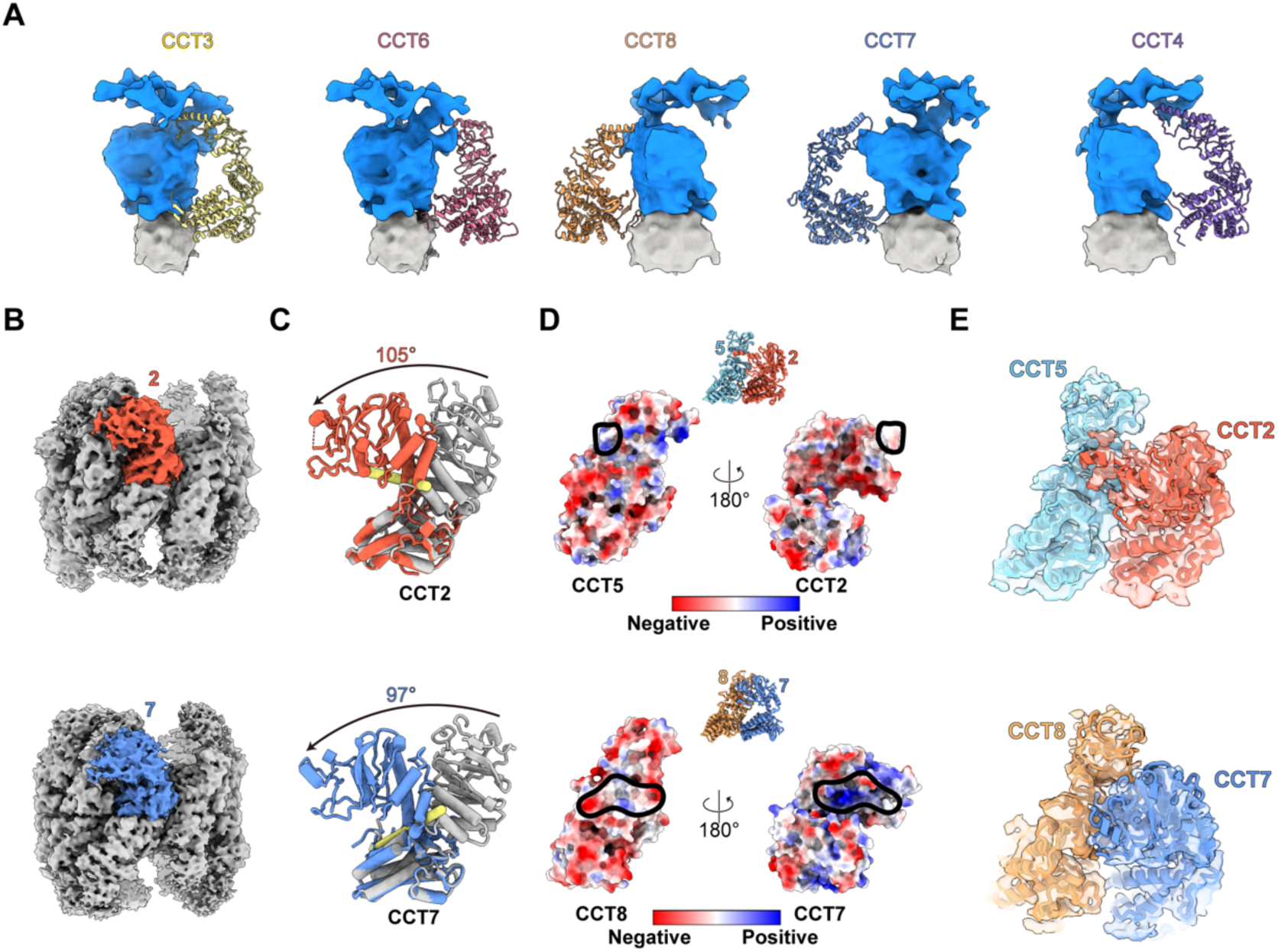
IFT172 engages the apo-state TRiC chamber through terminal tails and apical domain contacts. (A) Interaction analysis of the TRiC-IFT172 complex. The substrate density within the apo TRiC chamber contacts the disordered N- and C-terminal tails of TRiC subunits as well as the apical domains of multiple subunits. A continuous density extends from the chamber to the exterior, consistent with partial protrusion of IFT172 outside the cavity. (B) Structural representation of the Z-shaped conformations adopted by CCT2 and CCT7. (C) Measurement of bending angles for CCT2 and CCT7. Rotation axes (yellow) were automatically defined in ChimeraX. (D) Electrostatic surface potential analysis of the CCT2-CCT5 and CCT7-CCT8 subunit pairs, revealing stabilizing charge complementarity at the bent interface. (E) Model-to-map fitting of the Z-shaped CCT2-CCT5 and CCT7-CCT8 subunit pairs.

**Fig. S4.**
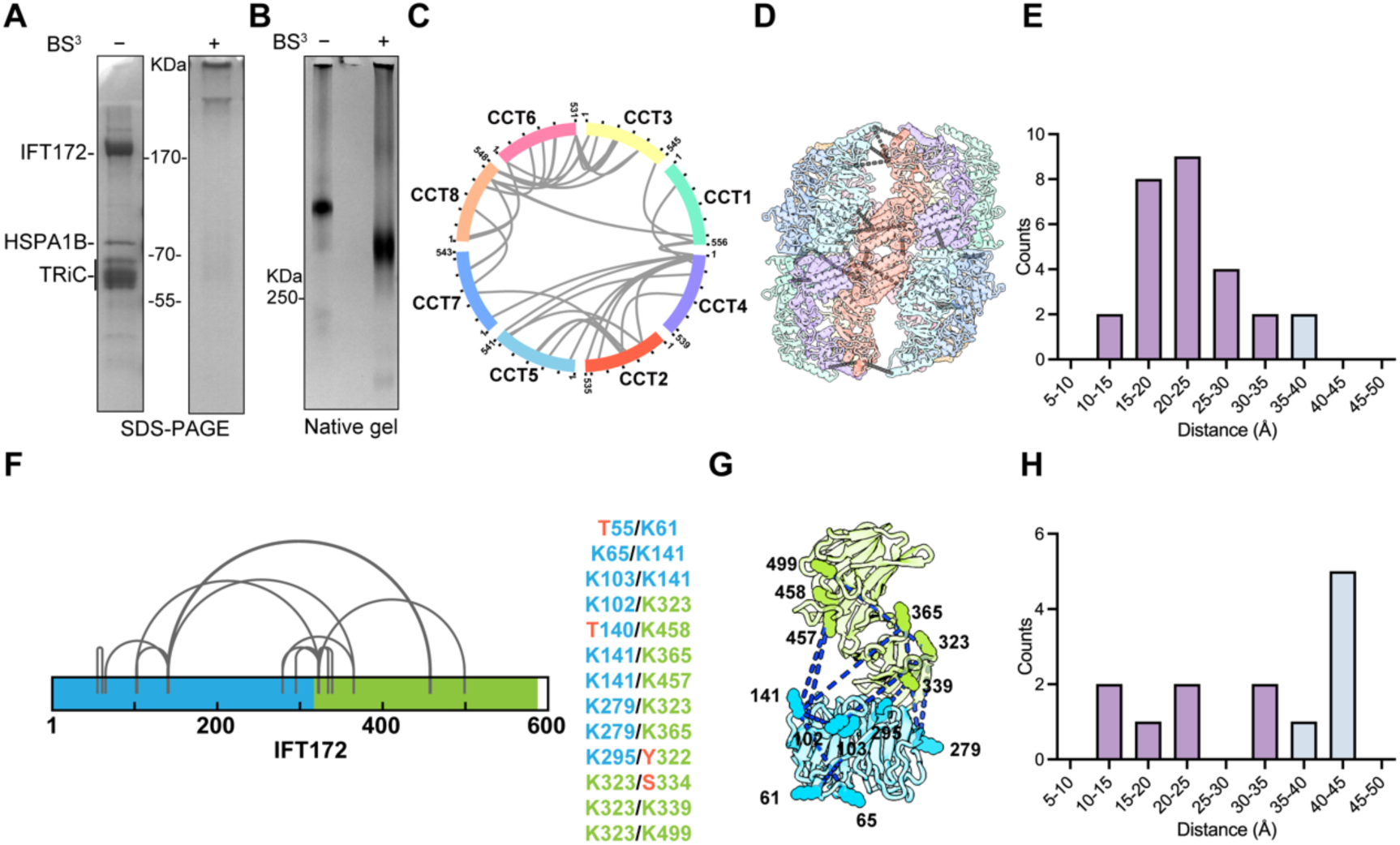
XL-MS defines domain-specific interactions between IFT172 and TRiC. (A) Coomassie-blue stained SDS-PAGE of the BS^3^ crosslinked TRiC-IFT172 complex. (B) Native PAGE (4%) showing a faster-migrating species after crosslinking, consistent with stabilized complex formation. (C) Circular plot of intra-TRiC inter-subunit crosslinks, consistent with the known subunit arrangement. (D) Mapping of TRiC crosslinks onto our apo-state TRiC model. (E) Distribution of Cα–Cα distances for intra-TRiC crosslinks; 92.6% fall within 35 Å, validating dataset quality. (F) Intra-molecular crosslinks within the IFT172 WD40 domain (WD40-1, blue; WD40-2, green). (G) Crosslinked lysine residues mapped onto the AlphaFold3-predicted IFT172 WD40 model. (H) Distribution of Cα–Cα distances for mapped IFT172 WD40 crosslinks; only 53.8% satisfy the 35 Å threshold, indicating a non-native conformation when bound to TRiC.

**Fig. S5.**
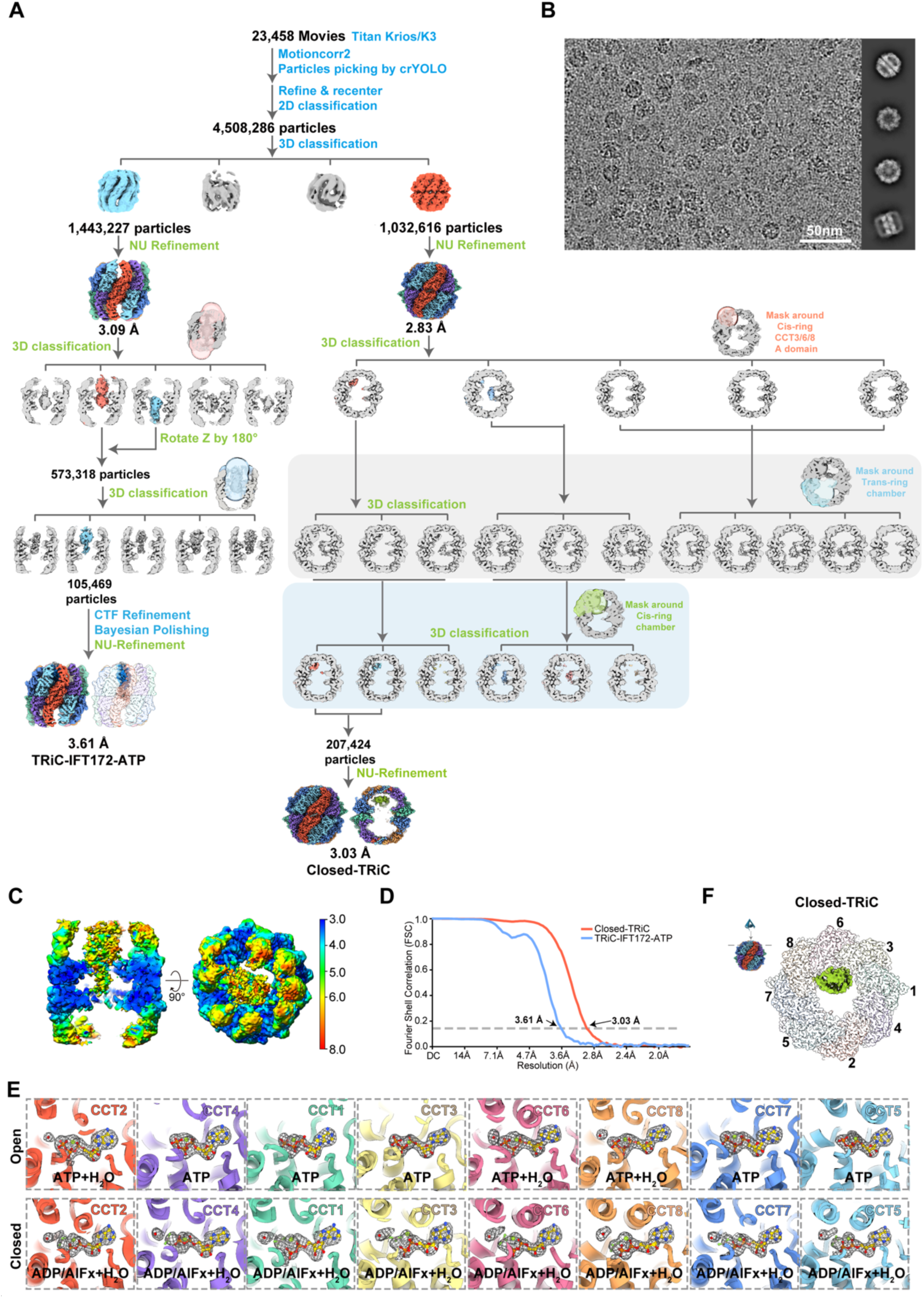
Cryo-EM analysis of the TRiC-IFT172 complex in the presence of ATP-AlFx. (A) Single-particle processing workflow for the TRiC-IFT172-ATP-AlFx dataset (without crosslinking). (B) Representative cryo-EM micrograph and reference-free 2D class averages. (C) Local resolution estimations for TRiC-IFT172-ATP map. (D) Fourier shell correlation (FSC) curves for the reconstructions (FSC= 0.143 criterion). (E) Model-to-map fitting of nucleotide-binding pockets in the open and closed TRiC states. In the open state, ATP is bound in all eight subunits, with putative attacking water molecules resolved in CCT2, CCT6 and CCT8. In the closed state, all subunits contain ADP-AlFx, with a coordinated Mg^2+^ ion (green ball) and an attacking water molecule (red sphere). (F) Additional density observed within the closed TRiC chamber under ATP-AlFx condition.

**Fig. S6.**
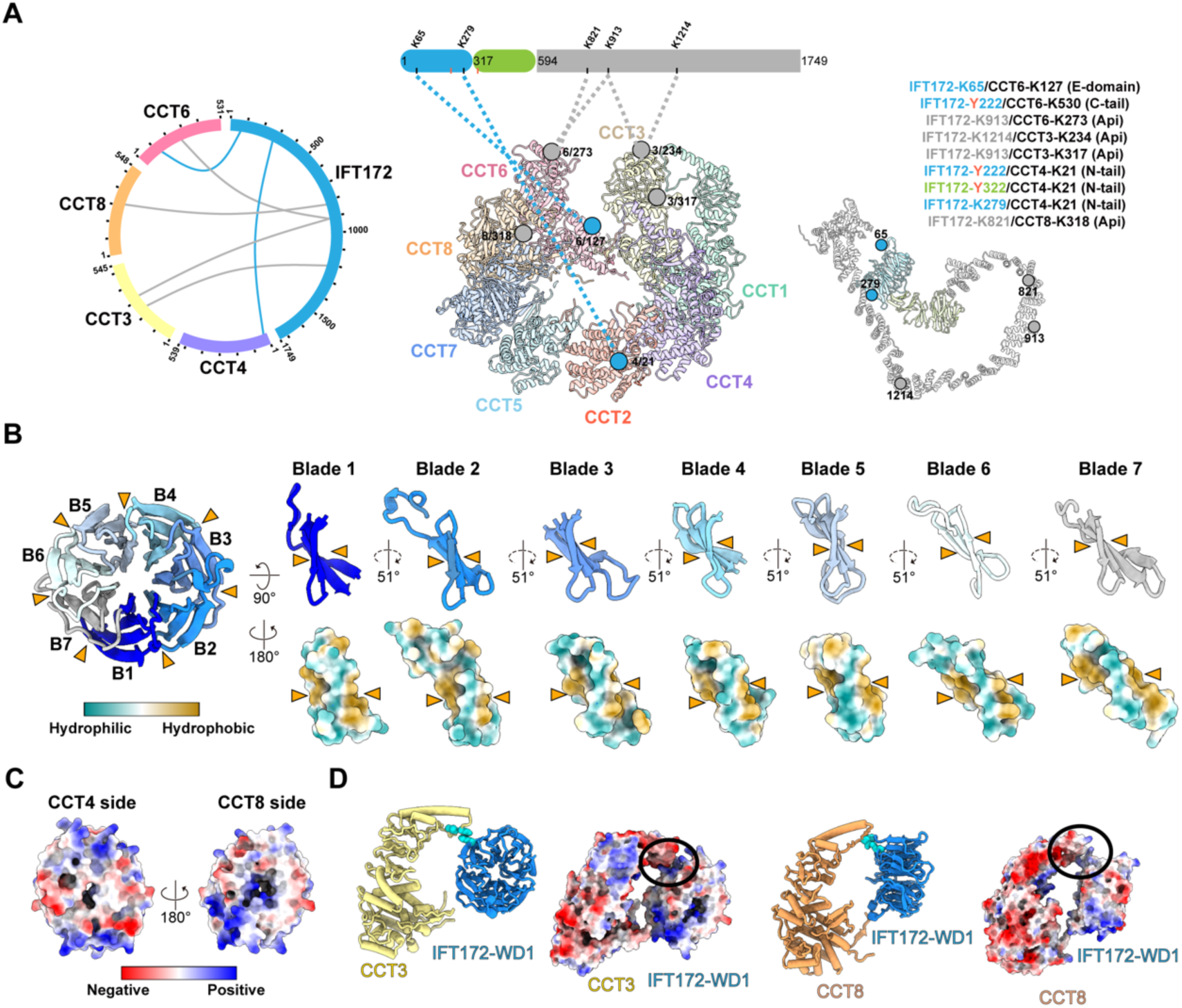
TRiC mediates folding of the IFT172 WD40-1 domain in the ATP-bound open state. (A) XL-MS analysis of the TRiC-IFT172 complex under ATP-AlFx condition. (B) Structural features of the IFT172 WD40-1 domain, illustrating the hydrophobic inter-blade packing that stabilizes the β-propeller fold. (C) Electrostatic surface potential of IFT172 WD40-1. (D) Electrostatic surface potential analysis indicating complementary surface properties between the apical domains of CCT3/CCT8 and WD40-1, which further stabilizes the folded domain within TRiC chamber.

**Fig. S7.**
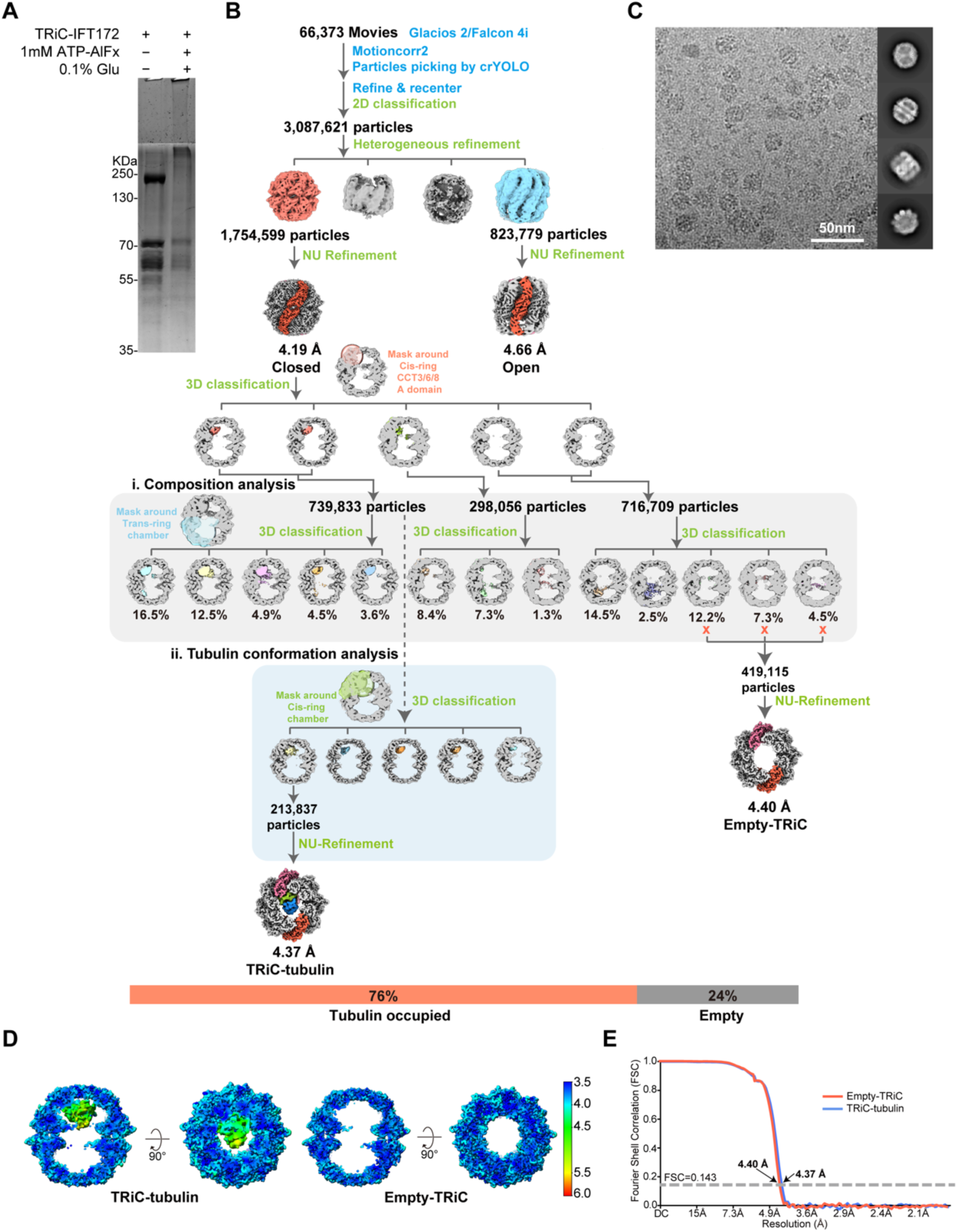
Cryo-EM analysis of cross-linked TRiC-IFT172 under ATP-AlFx condition reveals tubulin occupancy, but not IFT172, in closed chambers. (A) Coomassie blue-stained SDS-PAGE of TRiC-IFT172 treated with 1 mM ATP-AlFx and crosslinked with 0.1% glutaraldehyde. (B) Single-particle processing workflow for this dataset. Among closed-state TRiC particles, 24% contain an empty chamber, while the remaining 76% are occupied by endogenous tubulin in various folding states. (C) Representative cryo-EM micrograph and reference-free 2D class averages of the complex. (D) Local resolution estimations for the TRiC-tubulin and empty-TRiC maps. (E) Fourier shell correlation (FSC) curves for the empty closed TRiC map and tubulin-bound TRiC map (FSC=0.143 criterion).

**Table S1.**
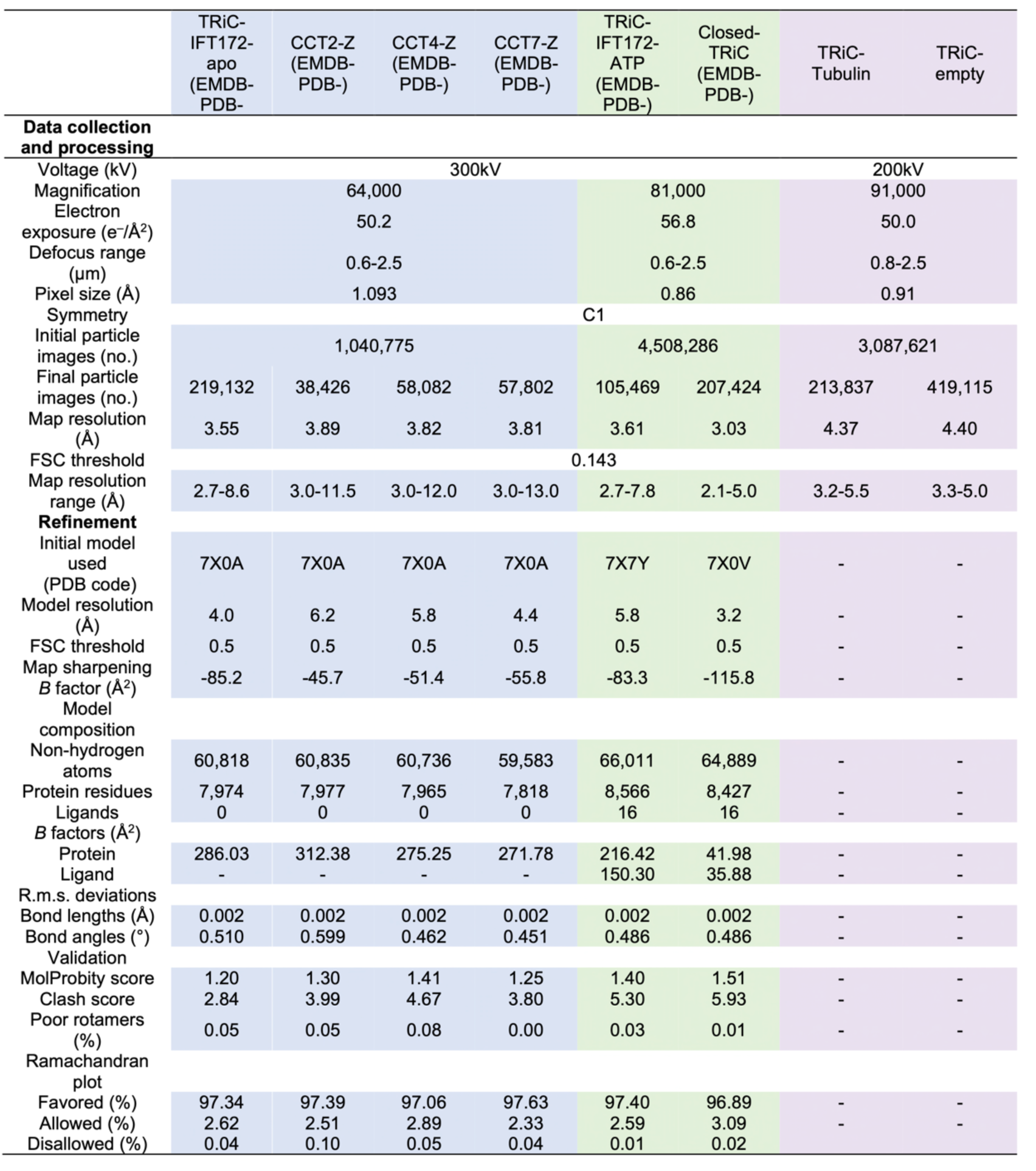
Cryo-EM data collection, processing, and model validation statistics.

**Table S2.**
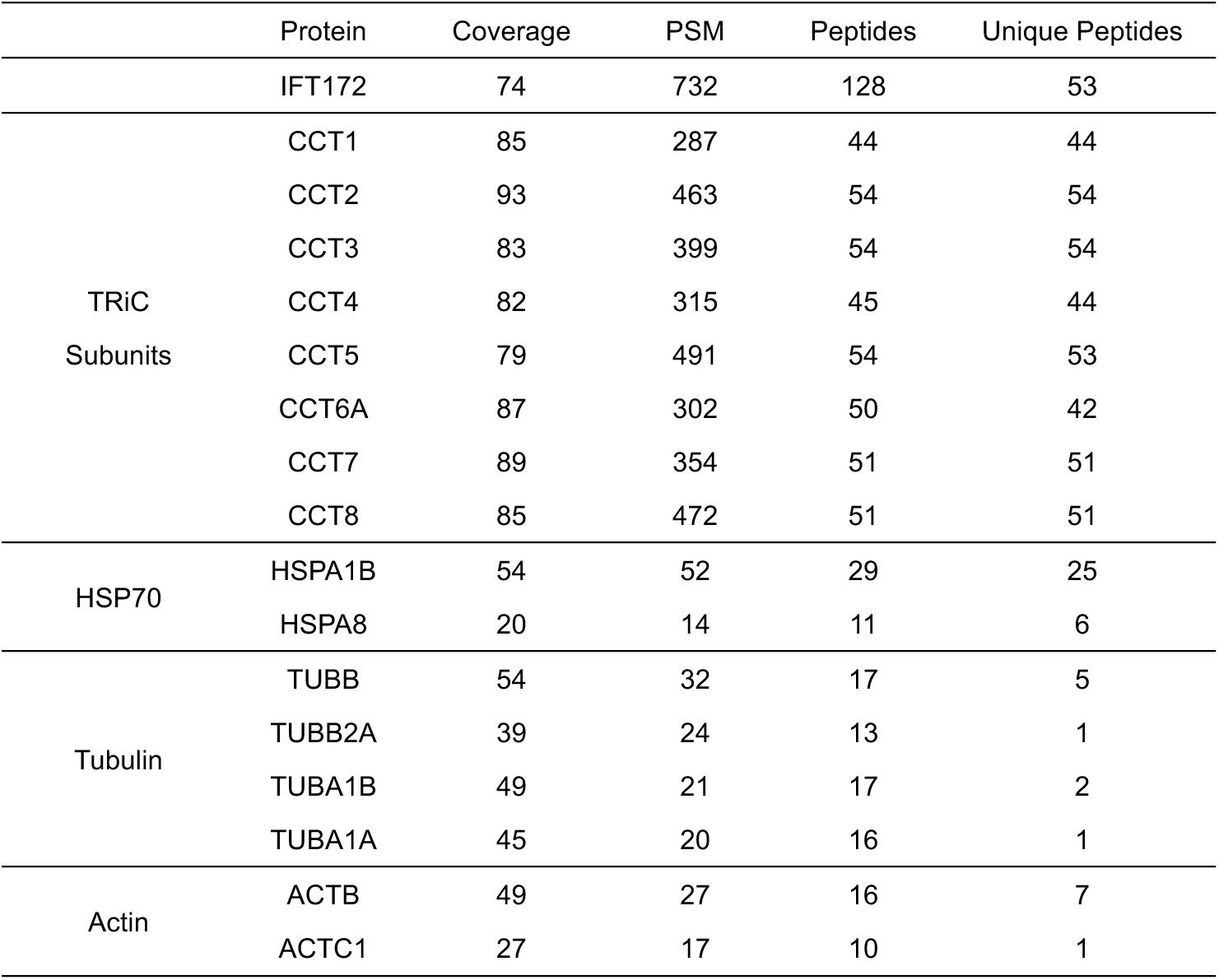
Mass spectroscopy analysis of the endogenously purified TRiC-IFT172 complex.

|  | Protein | Coverage | PSM | Peptides | Unique Peptides |
| --- | --- | --- | --- | --- | --- |
|  | IFT172 | 74 | 732 | 128 | 53 |
| TRiC<br>Subunits | CCT1 | 85 | 287 | 44 | 44 |
|  | CCT2 | 93 | 463 | 54 | 54 |
|  | CCT3 | 83 | 399 | 54 | 54 |
|  | CCT4 | 82 | 315 | 45 | 44 |
|  | CCT5 | 79 | 491 | 54 | 53 |
|  | CCT6A | 87 | 302 | 50 | 42 |
|  | CCT7 | 89 | 354 | 51 | 51 |
|  | CCT8 | 85 | 472 | 51 | 51 |
| HSP70 | HSPA1B | 54 | 52 | 29 | 25 |
|  | HSPA8 | 20 | 14 | 11 | 6 |
| Tubulin | TUBB | 54 | 32 | 17 | 5 |
|  | TUBB2A | 39 | 24 | 13 | 1 |
|  | TUBA1B | 49 | 21 | 17 | 2 |
|  | TUBA1A | 45 | 20 | 16 | 1 |
| Actin | ACTB | 49 | 27 | 16 | 7 |
|  | ACTC1 | 27 | 17 | 10 | 1 |

**Table S3.** Crosslinking mass spectroscopy analysis of the apo-state TRiC-IFT172 complex.

| Protein 1 (site)-Protein 2 (site) | Peptides | Best E-value | Spec count |
| --- | --- | --- | --- |
| IFT172(65)-CCT6(127) | KSYMVK(1)-IITEGFEEAAKEK(10) | 2.43E-22 | 15 |
| IFT172(66)-CCT6(127) | KSYMVK(2)-IITEGFEEAAKEK(10) | 5.87E-21 | 2 |
| IFT172(140)-CCT6(530) | LANTKTNK(4)-AGMSSLKG(7) | 5.78E-06 | 2 |
| IFT172(141)-CCT6(10) | LANTKTNK(5)-TLNPKAELVAR(5) | 1.09E-05 | 3 |
| IFT172(365)-CCT6(530) | ILGKER(4)-AGMSSLKG(7) | 2.25E-10 | 3 |
| IFT172(913)-CCT6(273) | NTASKYYPR(5)-KIIELEK(1) | 1.16E-04 | 3 |
| IFT172(141)-CCT3(21) | LANTKTNK(5)-KVQSGNINAAK(1) | 7.62E-06 | 6 |
| IFT172(295)-CCT3(21) | EIANLYTVTALAWKR(14)-KVQSGNINAAK(1) | 3.95E-12 | 2 |
| IFT172(1214)-CCT3(234) | GALEEKDFQK(6)-YIKNPR(3) | 3.26E-12 | 2 |
| IFT172(141)-CCT4(21) | LANTKTNK(5)-GKGAYQDR(2) | 5.04E-10 | 5 |
| IFT172(222)-CCT4(21) | IVAYGKEGHVLQTFDYSR(4)-GKGAYQDR(2) | 7.64E-11 | 2 |
| IFT172(224)-CCT4(21) | IVAYGKEGHVLQTFDYSR(6)-GKGAYQDR(2) | 6.63E-13 | 2 |
| IFT172(279)-CCT4(21) | SIWEEAKPK(7)-GKGAYQDR(2) | 4.09E-06 | 4 |
| IFT172(499)-CCT4(21) | VDWLELNETGHKLLFR(12)-GKGAYQDR(2) | 2.85E-10 | 3 |
| IFT172(632)-CCT4(21) | TLSKLALLEAR(3)-GKGAYQDR(2) | 1.91E-09 | 2 |
| IFT172(781)-CCT4(319) | AGLPAKAAR(6)-DALSDLALHFLNKMK(13) | 9.95E-09 | 3 |
| IFT172(140)-CCT2(522) | LANTKTNK(4)-VDNIIKAAPR(6) | 5.35E-06 | 2 |
| IFT172(141)-CCT2(522) | LANTKTNK(5)-VDNIIKAAPR(6) | 1.25E-07 | 5 |
| IFT172(323)-CCT2(522) | SIYKNK(4)-VDNIIKAAPR(6) | 7.21E-06 | 6 |
| IFT172(141)-HSPA1B(497) | LANTKTNK(5)-STGKANK(4) | 2.44E-04 | 2 |
| IFT172(821)-HSPA1B(345) | AGDLFEKIR(7)-IPKVQK(3) | 1.97E-03 | 2 |
| IFT172(821)-HSPA1B(497) | AGDLFEKIR(7)-STGKANK(4) | 3.84E-07 | 4 |
| IFT172(912)-HSPA1B(257) | NTASKYYPR(4)-DISQNK(6) | 6.74E-05 | 2 |
| IFT172(913)-HSPA1B(325) | NTASKYYPR(5)-DAKLDK(3) | 5.24E-09 | 5 |
| IFT172(913)-HSPA1B(246) | NTASKYYPR(5)-LVNHFVEEFKR(10) | 9.56E-13 | 2 |
| IFT172(913)-HSPA1B(507) | NTASKYYPR(5)-ITITNDKGR(7) | 4.23E-08 | 4 |
| IFT172(971)-HSPA1B(537) | PEDVSVLYITQAQEMEK(5)-<br>VSAKNALESYAFNMK(2) | 3.79E-07 | 2 |
| IFT172(1301)-HSPA1B(497) | AVDCYLKVR(7)-STGKANK(4) | 1.21E-08 | 2 |
| IFT172(913)-HSPA1B(345) | NTASKYYPR(5)-IPKVQK(3) | 3.24E-07 | 3 |
| IFT172(1382)-HSPA1B(345) | VAKELDPR(3)-IPKVQK(3) | 4.60E-08 | 2 |
| IFT172(1697)-HSPA1B(190) | NKIEFK(2)-TGKGER(3) | 5.55E-06 | 2 |
| IFT172(913)-HSPA1B(497) | NTASKYYPR(5)-STGKANK(4) | 1.03E-07 | 3 |
| IFT172(224)-HSPA1B(539) | IVAYGKEGHVLQTFDYSR(6)-<br>VSAKNALESYAFNMK(4) | 2.58E-13 | 2 |

**Table S4.** Crosslinking mass spectroscopy analysis of the ATP-AlFx presented TRiC-IFT172 complex.

| Protein 1 (site)-Protein 2 (site) | Peptides | Best E-value | Spec count |
| --- | --- | --- | --- |
| IFT172(65)-CCT6(127) | KSYMVK(1)-IITEGFEEAAKEK(10) | 3.58E-14 | 2 |
| IFT172(222)-CCT6(530) | IVAYGKEGHVLQTFDYSR(4)-AGMSSLKG(7) | 1.38E-08 | 2 |
| IFT172(913)-CCT6(273) | NTASKYYPR(5)-KIIELEK(1) | 3.51E-03 | 2 |
| IFT172(1214)-CCT3(234) | GALEEKDFQK(6)-YIKNPR(3) | 5.50E-15 | 4 |
| IFT172(913)-CCT3(317) | NTASKYYPR(5)-KTDNNR(1) | 3.22E-09 | 2 |
| IFT172(222)-CCT4(21) | IVAYGKEGHVLQTFDYSR(4)-GKGAYQDR(2) | 1.79E-14 | 2 |
| IFT172(322)-CCT4(21) | SIYKNK(3)-GKGAYQDR(2) | 3.67E-03 | 2 |
| IFT172(279)-CCT4(21) | SIWEEAKPK(7)-GKGAYQDR(2) | 7.53E-09 | 5 |
| IFT172(821)-CCT8(318) | AGDLFEKIR(7)-LNSKWDLR(4) | 1.43E-10 | 2 |

**Movie S1 3DVA of the apo-state TRiC-IFT172 complex.** Components 1 illustrate the coordinated intra-ring dynamics, while Component 2 captures the relative inter-ring movement.

**Movie S2. 3DVA of the ATP-bound TRiC-IFT172 complex.** Compared with the apo state (Movie S1), the ATP-bound complex displays markedly reduced conformational dynamics.

**Movie S3. Effect of TRiC RNAi on intraflagellar transport in *C. elegans*.** Neuron-specific RNAi-mediated knockdown of *cct-5* results in severe IFT deficiency, while *cct-8* knockdown leads to a significant reduction in IFT activity, compared with controls. See Fig. 5 for quantification.

